# Structural architecture of a dimeric paramyxovirus polymerase complex

**DOI:** 10.1101/2021.09.13.460081

**Authors:** Jin Xie, Li Wang, Guanglei Zhai, Daitze Wu, Zhaohu Lin, Manfu Wang, Xiaodong Yan, Lu Gao, Xinyi Huang, Rachel Fearns, Shuai Chen

## Abstract

Human parainfluenza virus type 3 (hPIV3), a member of non-segmented, negative-strand RNA viruses (nsNSVs), is the second most common cause of severe respiratory disease in pediatrics. The transcription and replication processes of nsNSVs are catalyzed by a multi-functional RNA-dependent RNA polymerase (RdRp) complex composed of the large protein (L) and the phosphoprotein (P). Previous studies have shown that the polymerase can adopt a dimeric form, however, the structure of the dimer and how it functions are not understood. Here we determined the cryo-EM structure of hPIV3 L-P complex at 2.7 Å with substantial structural details. A putative catalytic magnesium ion could be built in our structure, and structural comparison revealed atomic features conserved with other RNA viruses. Interactions identified between the two priming and intrusion loops and the connector domain potentially trigger the spatial movement of three C-terminal L domains for different steps of transcription and replication. Structural comparison with other nsNSV RdRps suggests common features of L-P binding. Furthermore, we report for the first time the structural basis of the L-L interaction in the partially modelled dimeric L-P structure, in which the connector domain of one L is positioned at the putative RNA template entry of the other L. Based on these findings, we propose a model by which L dimerization promotes efficient conversion of nascent RNA into a template.

---

The non-segmented negative-strand RNA viruses (nsNSVs) include numerous human pathogens, such as respiratory syncytial virus (RSV), human parainfluenza viruses (hPIVs) and rabies virus (RABV)^1^. HPIV3, one of four hPIV subtypes, is the second most common cause of severe viral respiratory diseases in infants and children^2,3^. The RNA genome of nsNSVs is packaged by the nucleoprotein (N) into a helical ribonucleoprotein (RNP) complex that acts as the template for genome transcription and replication by the RdRp complex^4–8^. The RdRp complex of hPIV3, like other nsNSV members, consists of the large protein (L) and the phosphoprotein (P)^9^ (Fig. 1a). L harbors five domains including the RdRp domain, polyribonucleotidyltransferase (PRNTase) domain, connector domain (CD), methyltransferase (MTase) domain, and the C-terminal domain (CTD), and possesses all enzymatic activities^10^. Consisting of an N-terminal domain (NTD), central oligomerization domain (OD) and C-terminal X domain (XD), the P protein performs critical regulatory functions through interactions with L and N proteins^11–14^. The past few years have witnessed progress in the structural studies of the nsNSV L-P polymerases from the rhabdoviruses, vesicular stomatitis virus (VSV)^10,15^ and RABV^16^, the pneumoviruses, RSV^17,18^ and human metapneumovirus (hMPV)^19^, and a paramyxovirus, canine PIV5^11^. However, the RNA transcription and replication mechanisms of nsNSV L-P are not well understood, and the structure of a human paramyxovirus polymerase remains elusive. In addition, the nsNSV polymerase has been reported to form a functional dimer^20–22^, but its structure has only been detected in low-resolution negative-stain electron microscopy^15,16,23,24^. Here, we report the cryo-electron microscopy (cryo-EM) structure of hPIV3 L-P polymerase at the highest resolution so far for nsNSVs. This study strengthens our understanding of priming, initiation and elongation mechanism of nsNSV polymerase, and reveals the structural conservation of L-P binding. More interestingly, the L-L interaction was directly observed and suggests a novel model of nsNSV genome replication.

**Fig. 1.**
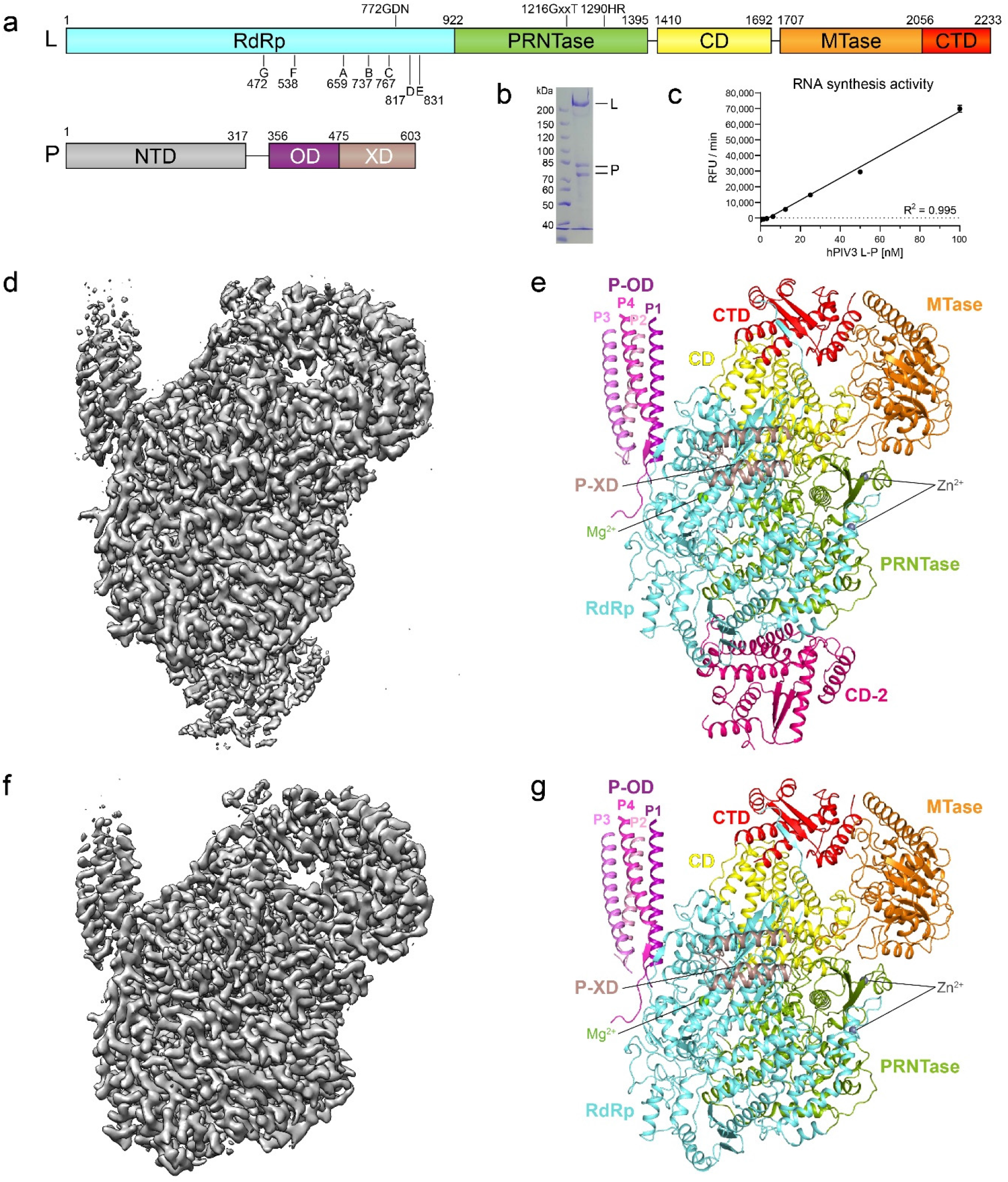
Overall structure of the hPIV3 L-P complex. **a,** Domain organization of L and P proteins. RdRp, aquamarine; PRNTase, green; CD, yellow; MTase, orange; CTD, red; P-NTD, grey, P-OD, purple; P-XD, darksalmon. The conserved motifs and residues required for functions are indicated. **b,** SDS-PAGE of the purified L-P complex. The second P band could represent a truncated product or alternative phosphorylated state. **c,** RNA synthesis activity was tested in a fluorescence-based primer extension assay. **d, e,** The cryo-EM map **(d)** and structure **(e)** of dimeric hPIV3 L-P with the CD domain (CD-2) of the second L. Domains are colored as depicted in **a** except for CD-2 in hot pink and four copies of P-OD domains in purple (P1), light pink (P2), violet (P3) and light magenta (P4), respectively. The magnesium ion at the RdRp active site and zinc ions in the PRNTase domain are shown as light green and grey spheres, respectively. **f, g,** The cryo-EM map **(f)** and structure **(g)** of monomeric hPIV3 L-P.

## 2.7 Å cryo-EM structure of hPIV3 L-P polymerase complex

Full-length human PIV3 L and P proteins (Fig. 1a) were co-expressed in Sf21 cells. The purified L-P complex (Fig. 1b) showed strong binding to an RNA duplex (Extended Data Fig. 3a) and its RdRp activity was verified by a fluorescence-based primer extension assay^25^ (Fig. 1c).

We employed single-particle cryo-EM to solve the hPIV3 L-P complex structure at a resolution up to 2.7 Å (Fig. 1d, e, Extended Data Fig. 1, Extended Data Table 1). This is the first nsNSV RdRp structure with a resolution higher than 3.0 Å, allowing us to build a more detailed atomic model. All five domains of L protein and four copies of OD domain and single XD domain of P protein were built into the density map. The overall structural architecture is similar to that of canine PIV5 L-P^11^. Interestingly, a large extra blob of electron density, not reported in previous L-P structures, was observed near the intact L protein and was successfully assigned as the CD domain of a second L protein (Fig. 1d, e, Extended Data Fig. 1). In addition to the partially modelled dimeric L-P structure, another main class of particles lacked density for the second CD domain and was reconstructed to a 3.3 Å resolution structure representing the monomeric L-P complex (Fig. 1f, g, Extended Data Fig. 1). Since these two structures show nearly the same arrangement except for the second CD domain, we used the 2.7 Å structure for further analysis.

## RdRp active site of hPIV3 L

Consistent with previously determined RNA polymerase structures^5,10,11,17,26–28^ (Extended Data Fig. 2), the RdRp domain of hPIV3 L-P complex folds into the canonical “right hand” fingers-palm-thumb subdomains, while the catalytic active site is composed of seven conserved motifs (A–G) (Fig. 2a, b). Motifs A-E are located in the palm subdomain, while motifs F and G are located in the finger subdomain. In order to further understand the RdRp active site of nsNSVs, we compared our structure to the RNA/Mg^2+^-bound RdRp structures of influenza B virus (FluB)^28^ and SARS-CoV-2^27^, as the representatives of segmented negative-strand RNA viruses (sNSVs) and positive-strand RNA viruses, respectively. The hPIV3 L RdRp domain shows similar structural architecture to that of FluB and SARS-CoV-2 (r.m.s.d of 2.93 Å over 349 Cα atoms and 3.43 Å over 319 Cα atoms, respectively) (Extended Data Fig. 2). Furthermore, motifs A–G could be well overlaid, and the proposed catalytic residues 772-GDN-774 of hPIV3 RdRp could be also superimposed with FluB (443-SDD-445) and SARS-CoV-2 (759-SDD-761) (Fig. 2c, d). One magnesium ion at the catalytic center could be built in our hPIV3 structure due to the well-resolved electron density (Fig. 2a–d, Extended Data Fig. 1f), which has not been reported in previous nsNSV L structures. The presumed catalytic Mg^2+^ is located at the similar position as one of the two Mg^2+^ present in FluB and SARS-CoV-2 structures, and coordinated by the side chain oxygen of the catalytic residue Asp773 and the main chain oxygen of motif A residue Leu664 (Fig. 2c, d). Similar to FluB^28^ and SARS-CoV-2^27^, hPIV3 L appears to possess the conserved motif F residues Arg552 and Lys543/Phe554 to stabilize the incoming nucleotide and the template strand RNA at the +1 site, respectively (Fig. 2c, d). In addition, motif G directs the template strand RNA into the active site with a conserved positively charged residue^27, 28^, Lys475 for hPIV3 (Fig. 2c, d). Consistent with the conserved nature of the RdRp domain, the broad-spectrum RdRp inhibitor remdesivir^27,29^ showed inhibition against hPIV3 in biochemical and virology assays (Extended Data Fig. 3b–e). Another RdRp inhibitor ALS-8112 against RSV and hMPV^30–32^ also showed activity against hPIV3 and VSV^31^.

**Fig. 2.**
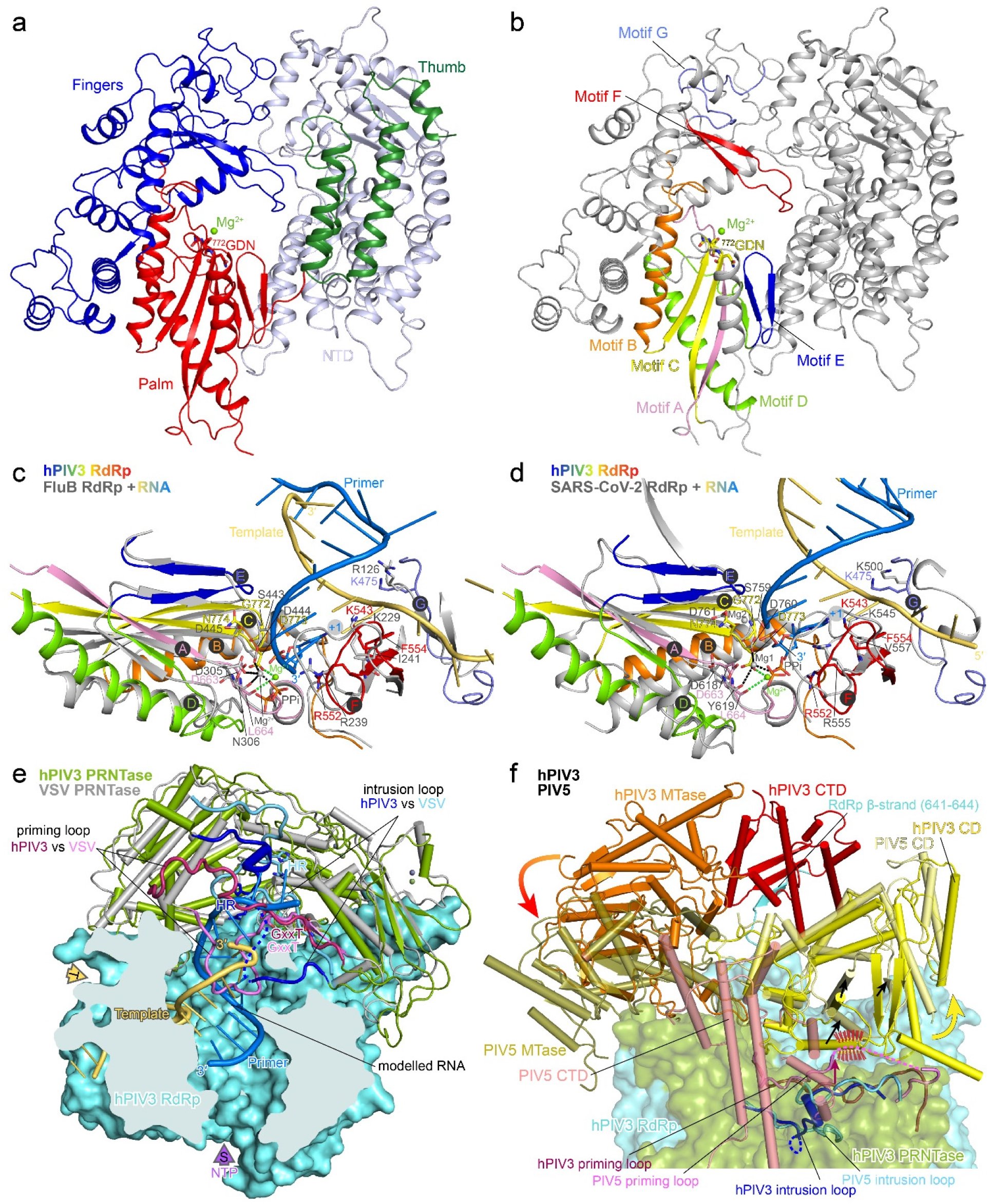
Structural architecture of the hPIV3 L protein. **a,** The RdRp domain is shown in ribbons, with the finger subdomain in blue, the thumb in dark green, the palm in red, and the N-terminal region (NTD) in blue-white. The catalytic residues 772-GDN-774 and the magnesium ion at the active site are shown as sticks and sphere, respectively. **b,** Motifs A–G of the RdRp domain are highlighted in rainbow colors with the same view as in **a**. **c, d,** RdRp active site. The RdRp domain of hPIV3 is superimposed on that of RNA/Mg^2+^-bound RdRp structures of FluB (PDB 6QCX) **(c)** and SARS-CoV-2 (PDB 7BV2) **(d)**. Motifs A–G and one magnesium ion in hPIV3 structure are depicted as in **b**. FluB and SARS-CoV-2 structures are colored in grey except for the template and primer strand RNAs in light orange and marine blue, respectively. The nucleotide at the +1 position of the primer strand and the pyrophosphate (PPi) are shown as sticks. The catalytic residues and critical conserved residues that interact with RNA are shown as sticks. **e,** The PRNTase domain (green) of hPIV3 L. A superposition of hPIV3 L with VSV L (PDB 6U1X) (grey) highlights the conformational differences of the putative priming loops and intrusion loops. The priming loops of hPIV3 and VSV are colored in claret and pink, respectively, while intrusion loops are in blue and cyan, respectively. The disordered region of hPIV3 intrusion loop is shown as a dotted line. The conserved GxxT motif and HR motif are labeled. The hPIV3 RdRp domain is shown as aquamarine surface and the RNA modelled from FluB structure above is shown as cartoon. **f,** Superposition of CD-MTase-CTD of hPIV3 (solid) and PIV5 (PDB 6V85) (transparent). RdRp-PRNTase of hPIV3 is shown as transparent surface. The disordered region of PIV5 priming loop is shown as a dotted line. Potential steric clash between PIV5 priming loop and the CD adopting the same conformation as hPIV3 is indicated by red dashes. Movements are indicated by arrows.

## HPIV3 L adopts a post-initiation conformation

The PRNTase domain of L is multifunctional, facilitating *de novo* initiation via a priming loop, capping the nascent viral mRNAs, and regulating RNA elongation^33–38^ (Fig. 2e, Extended Data Fig. 4). Compared to the relatively rigid structure of the core RdRp-PRNTase domains, the three C-terminal CD-MTase-CTD domains of L behave more dynamically^11,24^ (Extended Data Fig. 1c–e). The spatial position of the hPIV3 CD-MTase-CTD relative to RdRp-PRNTase is different from the reported nsNSV structures^10,11,15–19^ (Fig. 2f, Extended Data Fig. 5). A short β-strand extended from the hPIV3 RdRp domain forms a β-sheet, aiding the positioning of CTD. Comparison of the different structures suggests that they represent the RdRp at different stages of RNA synthesis.

In the reported VSV and RABV structures, the priming loop reaches towards the central cavity, representing a pre-initiation state^10,15,16^. In contrast, the hPIV3 priming loop (Val1243–Ser1259) retracts considerably from the central tunnel and props against CD, leading to the slight shift of CD and subsequent movement of MTase-CTD. Instead of the priming loop, an intrusion loop (Thr1280–Ser1305), containing the catalytic HR motif for capping, partially projects out into RdRp cavity, a feature also observed in PIV5 L^11^ (Fig. 2e, Extended Data Fig. 4a–d, 5a–d). Thus, both hPIV3 and PIV5 structures appear to be in a post-initiation state, but have several differences. In the hPIV3 structure, the distinct positioning of the HR motif towards the RNA cavity and the fact that the intrusion loop could accommodate extension of several base-pairs of RNA (Fig. 2e, Extended Data Fig. 4) might reflect a post-initiation state that occurs prior to mRNA cap addition, i.e. an RdRp at an earlier stage than that of PIV5^5,11^. The priming loop of PIV5 retracts further than that of hPIV3, resulting in the shift of PIV5 CD away from the cavity to avoid potential steric clash. The adjacent MTase-CTD module of PIV5 subsequently undergoes large conformational rearrangement positioning the MTase active site closer to the capping site (Fig. 2f, Extended Data Fig. 5c, d), which perhaps facilitates cap methylation^11^. Further structural differences are observed in the RSV and hMPV structures that adopt a non-initiation state^5,17–19^. Their priming and intrusion loops are fully retracted, possibly leading to a further shift of CD and more significant movement of MTase-CTD to expose the nascent RNA (Extended Data Fig. 5e, f). Consistent with this, their CD-MTase-CTD domains were not resolved in the EM maps^17–19^. In summary, comparison of our hPIV3 structure with other L-P structures revealed that the conformations of priming and intrusion loops are very likely linked to the movement of CD-MTase-CTD transitioning through different functional states.

## Unique and conserved L-P binding

More detailed L-P interactions are observed in our structure than in PIV5^11^ (Fig. 3a–h). Tetrameric OD domains of hPIV3 P constitute a long helical bundle bound to the RdRp domain of L with longer visible loops than PIV5 (Fig. 3i). Each of the four P monomers (P1–P4) adopts asymmetric conformations. Two proximal subunits P1 (Arg451–Arg470) and P4 (Leu464–Lys473) make extensive contacts with L (Fig. 3a–f). Residues Asn463–Glu469 of P4 are liberated from the long helix, and residues Lys465–Met467 form a typical antiparallel β-sheet with Gln387–Lys389 of L (Fig. 3e). In addition, the neighboring residue Phe390 along with Ile452 and Leu678 of L inserts into the exposed hydrophobic core composed of P1 and P4 (Fig. 3d). Similar β-sheet and hydrophobic interactions are also present in RSV and hMPV structures^17–19^. Among the three C-terminal α helices (named α1–α3 here) of hPIV3 P-XD, the α1 and its upstream loop, corresponding to the single α-helix at the C-terminus and the neighboring linker observed in RSV and hMPV L-P structures^17–19^, contribute to the majority of interactions between P-XD and L (Fig. 3g, h).

**Fig. 3.**
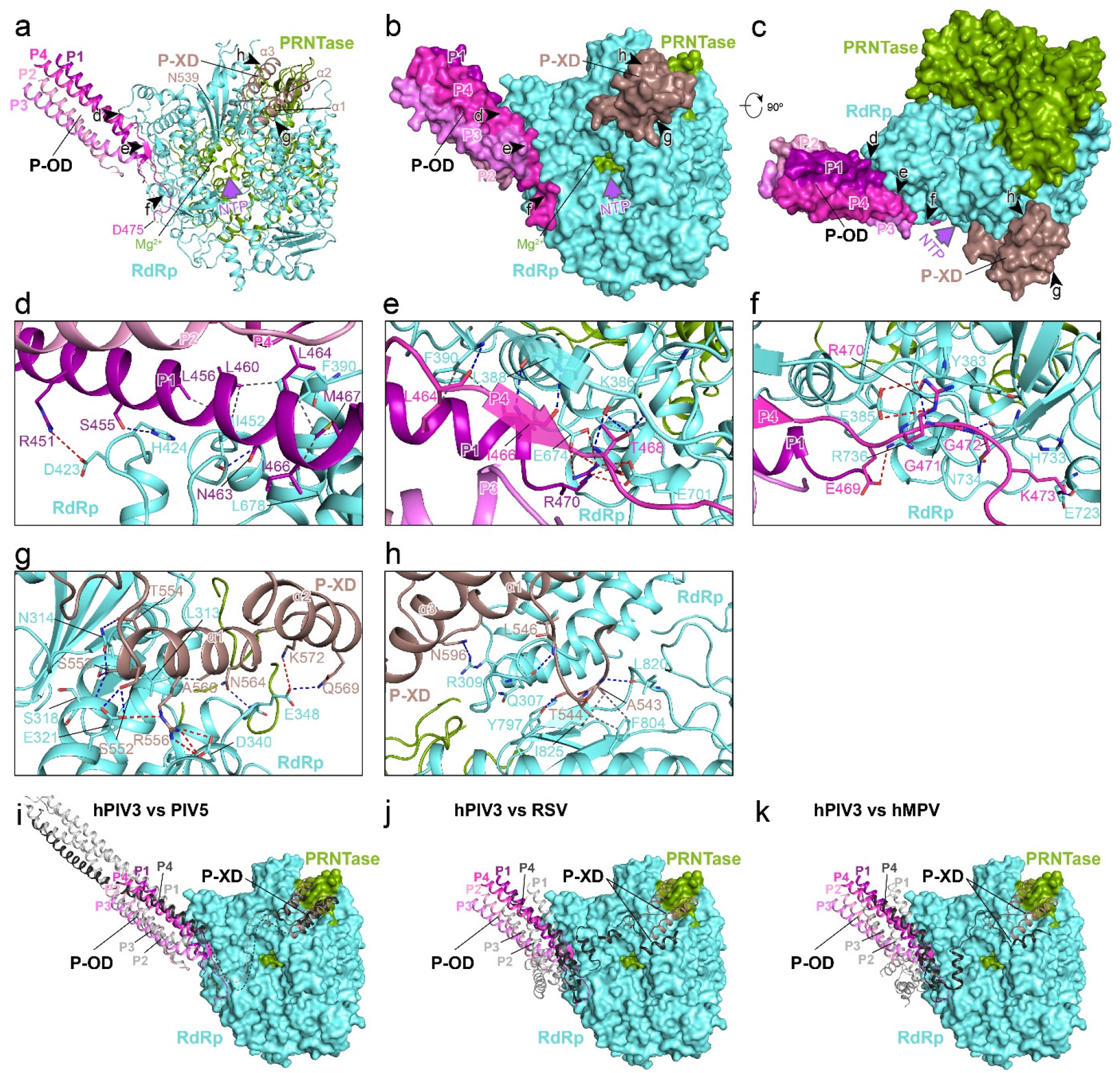
L-P interaction of hPIV3. **a,** Overview of OD and XD domains of P binding to L. Five arrows represent the views shown in **d**–**h**. L and P are shown as ribbons. The putative invisible linker between P1-OD and P-XD based on PIV5 L-P structure (PDB 6V85) is shown as a dotted line with the flanking residues labeled. The NTP entry is indicated by a purple arrow. **b,** The same view as in **a** with L and P shown as surface. **c,** Rotated view by 90° about the horizontal axis from **b**. **d**–**f,** Magnified view of the interaction between L and P-OD. Hydrogen bonds, salt bridges and representative hydrophobic interactions are indicated by dashed lines in blue, red and grey, respectively. **g, h,** Magnified view of the interaction between L and P-XD. **i**–**k,** Comparison of L-P binding between hPIV3 and PIV5 **(i),** RSV **(j),** hMPV **(k)**. Surfaces of hPIV3 RdRp and PRNTase are shown. HPIV3 L and P are colored as depicted in **a**. For PIV5, RSV (PDB 6PZK) and hMPV (PDB 6U5O), P4 that extends its C-terminal to the place roughly the same to P-XD of hPIV3 is colored in black, while another three P copies are colored in grey.

Most of the residues involved in the L-P interactions are highly conserved in the closely related paramyxoviruses such as caprine PIV3 (cPIV3), human PIV1 (hPIV1) and Sendai virus (SeV), while less conserved in the distantly related paramyxovirus PIV5 and other nsNSVs (Extended Data Figs. 8, 9). However, both OD and XD domains of hPIV3 and PIV5 P proteins adopt similar conformations and bind to the same approximate positions on L (Fig. 3i). Interestingly, RSV^17,18^ and hMPV^19^ also dock the tetrameric helical OD domains of P on the L surface like hPIV3 with some similar interaction features described as above, and extend their C-terminal helices of one monomer to roughly the same position as hPIV3 (Fig. 3j, k). In addition, linker density was observed between P4-OD and the C-terminal helices of P-XD in PIV5^11^, comparable to residues 163–210 of RSV P4. In summary, elements of L-P binding seem to be conserved between the *Paramyxoviridae* and *Pneumoviridae* families despite diversity in P sequence and structure.

## Structural definition of the L-L interaction

Genetic and biochemical evidence showed that L proteins of hPIV3^22^, hPIV2^39^ and SeV^21^ could form oligomers, and dimeric L-P complexes of RABV^16^ and VSV^15,23,24^ were observed under negative-stain EM, however the detailed structures and interactions remain unclear. Here, we obtained the partially modelled dimeric L-P structure of hPIV3 with the CD domain (named CD-2) of a second L protein clearly traced (Fig. 1d, e, Extended Data Fig. 1g). Some putative complete L-P dimers were also observed in 2D classification, consistent with the determined molecular weight of the complex (Extended Data Fig. 6).

CD-2 contacts the RdRp and PRNTase domains of the visible full L, with a 2452 Å^2^ total buried interface composed of a number of electrostatic and hydrophobic interactions (Fig. 4a, b). The charged residues Arg1509, Asp1510, Asp1576 and Glu1620 on CD-2 form salt bridges with Glu1112, Arg1101, Arg1116 and Arg1119 on a long α-helix of PRNTase (Fig. 4c–e). Some hydrogen bonds and hydrophobic interactions are formed between residues of CD-2 on helices α5 and α9 along with its upstream loop and residues from both RdRp and PRNTase domains of the intact L (Fig. 4c–e). In addition, Trp1513 of CD-2 forms cation–*π* interaction with Arg1101 of PRNTase (Fig. 4d). CD-2 has a similar overall structure as the CD domain of the intact L, except for the two loops that contribute to the L-L interface and the bent helix α2 of CD-2 (Extended Data Fig. 7a). It reveals that the two CD domains can bind to distinct interfaces on RdRp and PRNTase domains (Extended Data Fig. 7b), thus acting as both intramolecular and intermolecular connectors. Although the CD domain has no known enzymatic activity, it cannot tolerate in-frame insertions and domain exchanges between the substrains^40^. The residues involved in the L-L interactions appear to be conserved only among the closely related paramyxoviruses (Extended Data Fig. 8). However, despite a low sequence identity of 21%, hPIV3 and PIV5^11^ CD domains share very similar structures, and modelling of PIV5 L-P dimer showed some potential comparable L-L interactions. Taken together, the L-L dimerization interface may be conserved across paramyxoviruses and possibly other nsNSVs.

**Fig. 4.**
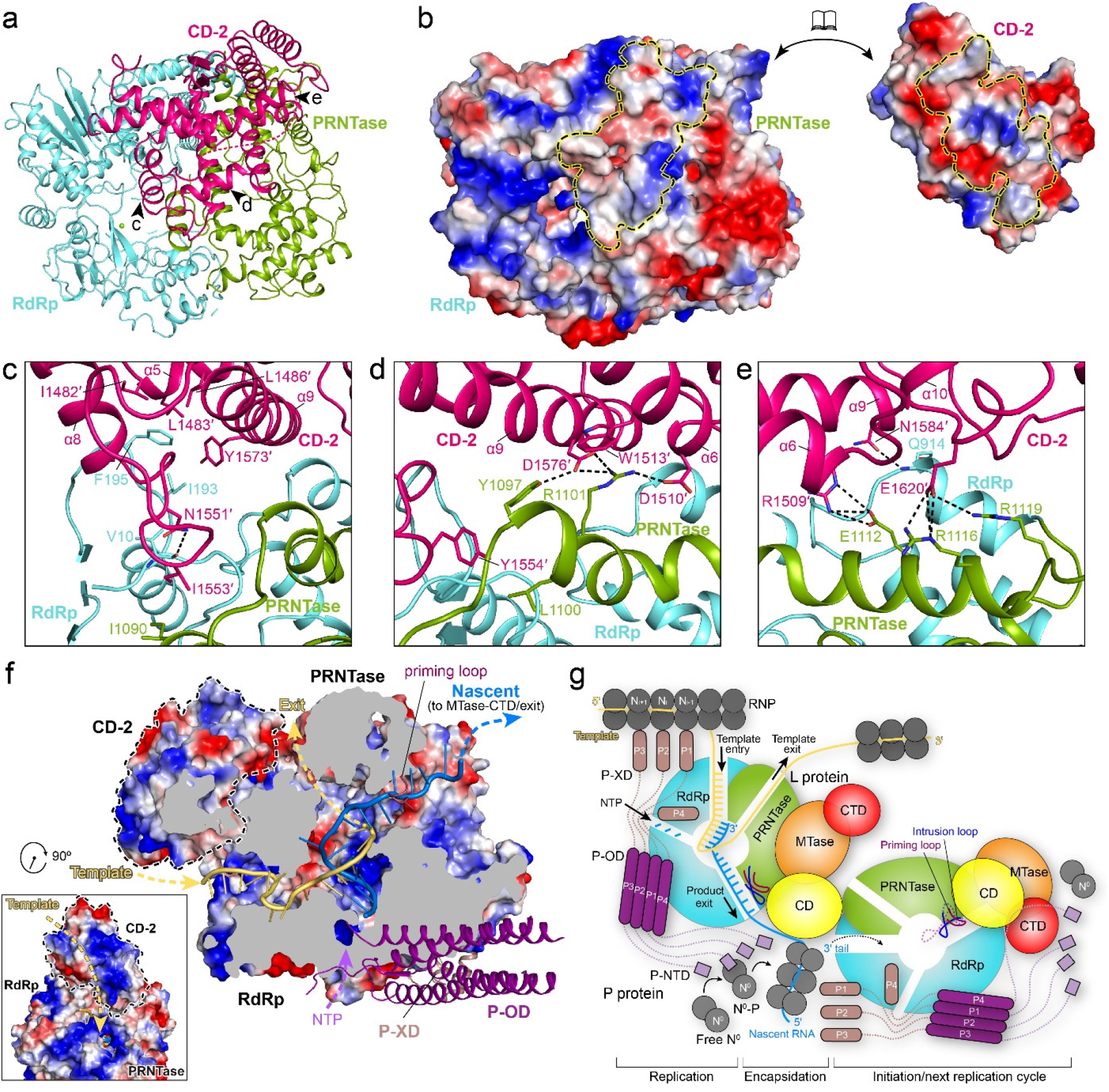
L-L interaction and dimeric L-P polymerase model. **a,** Overview of the CD domain (CD-2) of the second L binding to the RdRp and PRNTase domains of the intact L. Three arrows represent the views shown in **c**–**e**. **b,** The surfaces of CD-2 and RdRp-PRNTase are colored by electrostatic potential from red to blue (negative to positive, respectively). The interface of each is indicated by a dashed circle. **c**–**e,** Magnified view of the interaction between CD-2 and RdRp-PRNTase. The polar interactions are indicated by dashed lines. **f,** Cut-out view of L RdRp and PRNTase domains bound with CD-2 showing electrostatic surfaces and paths followed by the template (light orange) and nascent (marine blue) RNA strands modelled from FluB RdRp structure (PDB 6QCX). NTP entry, template RNA entry and exit, and nascent RNA exit tunnels are indicated by dashed arrows. The part that affects the nascent RNA exit in the priming loop is omitted from the electrostatic surfaces and shown as ribbons. Rotated view by 90° about the z axis is shown at the left bottom. **g,** The dimeric L-P polymerase model in RNA replication. The putative priming and intrusion loops in pre-initiation state are indicated by dashed transparent lines.

## Model for dimeric L-P polymerase

Recently, structures of both symmetric and asymmetric RNA polymerase dimers were reported for influenza viruses, enhancing our understanding of the role of RdRp oligomerization in RNA replication of sNSVs^5,41–43^. Here, the hPIV3 L-P dimer we captured appears to be asymmetric, otherwise serious steric clashes would be introduced between two L-P monomers. The fact that the rest of the second L-P was not visible in our reconstructed structure suggests that the second L-P monomer may adopt a non-initiation state similar to RSV and hMPV structures^17–19^ with dramatic movement flanking CD (Extended Data Fig. 5e, f). Interestingly, CD-2 is positioned to the putative template entry of the neighboring L, and has a conserved positively charged inner surface connected to the RNA tunnel (Fig. 4f and Extended Data Figs. 7d, 8). A similar basic path connecting the product exit channel of the replicating polymerase to the template entry on the encapsidating polymerase was also observed in the asymmetric dimer of influenza virus^43^. Previous studies have shown that inactive SeV L variants with mutations in different domains could complement each other to restore mRNA transcription and genome replication albeit at a low level, and complementation for leader RNA synthesis is efficient, indicating that the dimerization or oligomerization of L-P polymerase might be required for transcription and replication^21^. Based on these findings and the captured asymmetric dimer of hPIV3 L-P, we proposed a dimeric L-P polymerase model for nsNSV RNA replication (Fig. 4g), although we cannot exclude the possibility that nsNSV transcription also needs dimeric polymerase. P-XD first binds to N protein in the RNP complex and scans along the genome^6^. The 3′ terminus of the template RNA is loaded onto the central RNA tunnel to initiate RNA synthesis. The priming loop then retracts into the cavity of PRNTase, together with the intrusion loop, pushing away the CD domain and triggering dramatic movement of the adjacent C-terminal MTase-CTD. The opening of the product exit allows nascent RNA to travel along the positively charged surface of the CD domain. The NTD of P located near CD and CTD of L^15,16^, recruits the RNA-free N protein (N^0^) via its N-terminal tail^44^ to encapsidate the nascent RNA. After the replication of template RNA and encapsidation into a helical nucleocapsid, the 3′ terminus of the product emerges to be recognized by the second L-P, which transits from the initiation state to the replication state to start a new cycle.

## Methods

### Cloning and expression

Human PIV3 L (Gene ID: 911960) and P (Gene ID: 911956) genes codon-optimized for expression in *Spodoptera frugiperda* 21 (Sf21) cells, were chemically synthesized and cloned into pFastBac Dual transfer vector (Thermo Fisher Scientific). HPIV3 L (M1–D2233) with a C-terminal FLAG tag (DYKDDDDK) and 25 additional N-terminal residues MISNQQSDNGQKENIKNLGAKRARK was inserted downstream of the polyhedrin promoter and P (M1–Q603) with a C-terminal His_6_ tag was inserted downstream of the p10 promoter in pFastBac Dual. Recombinant bacmid containing hPIV3 L-P genes was generated and isolated following the instruction manual of the bac-to-bac baculovirus expression system (Thermo Fisher Scientific). Viral stocks generated from purified bacmids were amplified and used for protein expression. 2.4 L of Sf21 cells were infected with amplified viruses to co-express hPIV3 L and P proteins at 27°C for 48 h.

### Protein purification of hPIV3 L-P complex

The pellet of Sf21 cells expressing the hPIV3 L-P complex was resuspended through high-pressure homogenizer in the lysis buffer (50 mM Tris, pH 7.5, 500 mM NaCl, 10% glycerol, 6 mM MgSO_4_, 1 mM dithiothreitol (DTT), 1% Triton X-100) supplemented with cOmplete™, EDTA-free protease inhibitor cocktail (Roche). After high-speed centrifugation at 5,0000× g for 60 min at 4°C, the supernatant containing the target proteins was loaded onto anti-FLAG G1 affinity resin (Genscript). The resin was washed using buffer A (20 mM Tris, pH 7.5, 300 mM NaCl, 5% glycerol, 6 mM MgSO_4_, 1 mM DTT), and the bound proteins were eluted using 0.2 mg/ml FLAG peptide in buffer A. The eluted proteins were further purified using a size-exclusion column (Superose 6 Increase 10/300 GL, GE healthcare) equilibrated with buffer B (20 mM Tris, pH 7.5, 300 mM NaCl, 1% glycerol, 6 mM MgSO_4_, 1 mM DTT). The peak fractions were collected and analyzed by sodium dodecyl sulphate–polyacrylamide gel electrophoresis (SDS-PAGE). Fractions that contained the L-P complex were combined and concentrated to 1.2 mg/ml. The final sample was flash-frozen, and stored at −80 °C. The protein homogeneity was characterized by negative-stain EM. Molecular weight (MW) of the purified hPIV3 L-P complex was determined by analytical size-exclusion chromatography using a Superdex 200 Increase 5/150 GL column (GE healthcare) with buffer B. The standard molecular markers (cytiva, 28403842) were used for calibration. In addition, size-exclusion chromatography with multi-angle light scattering (SEC-MALS) was also used for MW determination of the complex. The purified hPIV3 L-P complex was loaded onto a XBridge™ Protein BEH SEC column (200 Å, 3.5 μm, 7.8 × 150 mm) (Waters, p/n: 176003595) using Breeze 2 high-performance liquid chromatography system (Waters) coupled with DAWN® MALS detector (WYATT-920-H2) at 0.5 ml/min. BSA was used as a control.

### Fluorescence-based primer extension assay

For the determination of the hPIV3 L-P RdRp activity, a real-time primer elongation assay was established utilizing the fluorescence dye SYTO9 (Thermo Fisher Scientific, S34854), which binds to double-strand RNA but not single-strand RNA^25,45^. A 40-nt RNA oligonucleotide with the sequence of 5′-UUUUUUUUUUUUUUUUUUUUUAGCUUCUUAGGAGAAUGAC-3′ was used as the template strand, and a 20-nt RNA oligonucleotide with the sequence of 5′-GUCAUUCUCCUAAGAAGCUA-3′ was used as the primer strand. To prepare the RNA duplex, 100 μM of both oligonucleotides were mixed at equal volume in the annealing buffer (10 mM Tris-HCl, pH 8.0, 25 mM NaCl, 2.5 mM EDTA, and 0.57 U/μl RiboLock RNase inhibitor (Thermofisher, EO0384)), denatured by heating to 94°C for 5 minutes, and then slowly cooled to room temperature. The primer extension assay was performed in a 384-well plate (PerkinElmer, 6008269). The reaction buffer contained 20mM Tris-HCl, pH 8.0, 10 mM NaCl, 10 mM KCl, 6 mM MgCl_2_, 0.01% TritonX-100, 1 mM DTT, 5% glycerol, 0.57 U/μl RiboLock RNase inhibitor. The purified hPIV3 L-P complex with a final concentration ranging from 0 to 100 nM was added into the reaction mixture containing 0.2 μM RNA duplex, 2 mM ATP and 0.125 μM SYTO9, to initiate the RNA elongation. The ENVISION (PerkinElmer) was employed to record the fluorescence signal using excitation wavelength and emission wavelength at 485 and 535 nm, respectively in real-time for 60 minutes at 25°C. 100 nM of hPIV3 L-P complex was adopted for the following assays. In the competition assay, the ATP analog, RDV-TP that is the triphosphate form of remdesivir at 200 μM was incorporated to compete against variant concentrations of ATP from 0 to 2 mM. For the IC_50_ value determination of RDV-TP against hPIV3 L-P complex, 0–0.5 mM of RDV-TP was added to the reaction mixture. Other conditions were the same as above.

### Fluorescence-based plaque reduction assay

The recombinant virus strain of hPIV3 that expresses enhanced green fluorescent protein (hPIV3-GFP) was purchased from ViraTree (P323). The human laryngeal cancer cell line (HEp2) was purchased from ATCC (CCL-23) and cultured in DMEM/F12 medium (Gibco, 11320033) with 10% heat-inactivated fetal bovine serum (FBS) and 1% penicillin/streptomycin. 10,000 cells/well of HEp2 cells were seeded in 96-well plates for 24 hours followed by infection of hPIV3-GFP at 0.02 M.O.I. in DMEM/F12 medium containing 2% heat-inactivated FBS and 1% penicillin/streptomycin at 37°C with 5% CO_2_. Remdesivir with a concentration ranging from 1 nM to 10 μM was added before viral infection and the final concentration of DMSO was 1%. 3 days after viral inoculation, plaques were detected in AID iSpot EliSpot FluoroSpot reader (Autoimmun Diagnostika GmbH, Germany) and counted using the software Algorithm C. Plaque reduction titers were calculated by regression analysis of the dilution of remdesivir compared to control wells incubated with 1% DMSO.

### Surface plasmon resonance (SPR) assay

The binding affinity between the purified hPIV3 L-P complex protein and a RNA duplex were measured at room temperature using a Biacore S200 system (GE Healthcare) with a SA chip (Cytiva, BR100531). The RNA duplex with the same sequence as that used in the primer extension assay except for 5′-biotin at the primer strand was immobilized on the chip. A blank channel of the chip was used as the negative control. The running buffer contained 10 mM HEPES, pH 7.4, 150 mM NaCl, 0.05% surfactant P20 and 6 mM MgCl_2_. Proteins were serially diluted to 1.95~250 nM using the running buffer and then loaded to flow through the chip surface. After binding and dissociation of each sample, the chip was regenerated using 50 mM NaOH with a short contact time of 5s. The sensorgrams were analyzed using the Biacore S200 Evaluation Software (version1.1) with 1:1 kinetics binding model.

### Cryo-EM sample preparation and data collection

An aliquot of 3.5 μL of hPIV3 L-P complex at 0.8 mg/ml was applied to a freshly glow-discharged Quantifoil R1.2/1.3 300-mesh grid. The sample was immediately blotted at 4°C and 100% relative humidity, then plunge-frozen in liquid ethane using Vitrobot Mark IV (Thermo Fisher Scientific, USA), and finally stored in liquid nitrogen. Cryo-EM data were collected on a Titan Krios microscope (Thermo Fisher Scientific, USA) under 300 kV, equipped with a K3 Summit direct electron detector (Gatan, USA). Movie stacks were automatically recorded using AutoEMation^46^ in the super-resolution mode at a nominal magnification of 81,000×, corresponding to a physical pixel size of 1.087 Å. The defocus was set from −1.5 to −2.0 μm. A total exposure dose of 50 e^-^/Å^2^ was fractionated into 32 frames for each movie stack. Finally, we obtained one cryo-EM dataset of hPIV3 L-P complex including a total number of 5,800 movie stacks.

### Cryo-EM image processing

A flow chart of cryo-EM data processing is shown in Extended Data Fig. 1. All dose-fractionated movie stacks were motion-corrected with RELION’s own motion-correction implementation^47^, yielding micrographs of 1.087 Å pixel size. After contrast transfer function (CTF) estimation using cryoSPARC^48^ Patch-CTF, a total of 5693 micrographs were selected for subsequent processing. To generate templates for automatic particle picking, 642 micrographs were selected, and 589,198 particles were auto-picked using cryoSPARC’s blob picker and extracted with a box size of 128 pixels after binning. After 2D classification, 100,387 particles were selected for 3D classification using Ab-Initio reconstruction in cryoSPARC, and three classes were generated as the initial reference models. Then 50 2D-templates were projected from the model with clear structural features. For the dataset of hPIV3 L-P complex, 4,511 micrographs were selected based on fitted resolution better than 4 Å, and a total of 3,288,374 particles were picked using templates generated previously and extracted with a box size of 150 pixels after binning. 636,487 particles were selected after two rounds of 2D classification based on the complex integrity. Then heterogeneous refinement was performed using previously generated Ab-Initio models. A subset of 373,483 particles from the class showing clear structural features was selected and re-extracted with a box size of 360 pixels without binning, and the resolution reached 3.3 Å after homogeneous refinement. For further classification, the full complex model and two erased models were used as 3D volume templates for heterogeneous refinement. The two classes of high quality (class 1 and 2, with and without a large extra blob of electron density, respectively) were subjected to homogeneous refinement, local refinement and non-uniform refinement using cryoSPARC. Then, CTF refinement and Bayesian polishing refinement were performed in RELION^47^. Finally, we obtained the 2.7 and 3.3 Å cryo-EM density maps for class 1 and class 2, respectively. In addition, several 2D classes generated from 46, 722 particles had a bigger size and appeared to be dimeric complex, but we failed to reconstruct further for these classes.

### Model building and refinement

The canine PIV5 L-P structure (PDB entry: 6V85)^11^ was used to guide the building of the atomic model of the hPIV3 L-P class 1 and 2. The complex of RdRp-PRNTase domains of L and four copies of P-OD and single P-XD of PIV5 was placed and rigid-body fitted well into the class 1 cryo-EM map using UCSF Chimera^49^. The CD and MTase-CTD domains of L were rigid-body docked separately with some rotation. Manual model building was carried out using Coot^50^ and refinement of the coordinates was performed using phenix.real_space_refine^51^. One magnesium ion at the catalytic center could be built due to its well-resolved electron density and close distance (about 2 Å) to the side chain oxygen of L residue Asp773. For the large extra blob of electron density in class 1 map, the main chains were firstly traced based on the excellent continuity of the electron density and obvious secondary structure feature, and bulky side chains visible in the density were utilized to determine the correct register of residues. Then the rough model was superimposed to the already built hPIV3 L-P structure and the result showed that it shared very high structural similarity to the CD domain of hPIV3 L. Based on the critical hint, this region was unambiguously built into the cryo-EM map and assigned as the CD domain of the second L protein in class 1 model. The class 2 model was then built based on the class 1 model. The final hPIV3 L-P model of class 1 comprises: the full L protein residues except for the N-terminal 7 residues, Tyr611–Lys637, Leu1292–Met1299, Ile1693–Asp1706, Thr1745–Thr1762 and Thr2095–Lys2113; the P protein residues with four copies of OD domains (Asp435–Gly471, Ala434–Met467, Asp435–Gly472 and Asp435–Asp475 of subunits P1, P2, P3 and P4, respectively) and single XD domain (Asn539–Gln603); and the CD domain of second L protein residues Asp1450–Leu1483 and Ile1495–Ile1712. The final hPIV3 L-P model of class 2 comprises nearly the same residues as class 1 except for the second CD domain. MolProbity^52^ was used to validate the geometries of the final models and the statistics are given in Extended Data Table 1.

### Figure preparation

UCSF Chimera^49^ and Pymol (Schrödinger LLC) were used for structure visualization and figure generation. Multiple sequence alignment was performed with MultAlin (http://multalin.toulouse.inra.fr/multalin/multalin.html) and the alignment results were displayed with ESPript^53^. The buried interface was calculated by PDBePISA^54^.

## Data availability

Structure coordinates are available from the RCSB Protein Data Bank (PDB) under accession codes 7V70 (hPIV3 L-P class 1, dimeric form) and 7V71 (hPIV3 L-P class 2, monomeric form), and the electron density map from the Electron Microscopy Data Bank (EMDB) under accession codes EMD-31755 (hPIV3 L-P class 1, dimeric form) and EMD-31756 (hPIV3 L-P class 2, monomeric form). All other data generated or analyzed in this work are available from the corresponding authors upon reasonable request.

## Acknowledgements

We appreciate the cryo-EM facility at Westlake University for the assistance in cryo-EM data acquisition. We would like to thank Zhanyu Ding from Shanghai YueXin Life-science Infomation Technology Co. Ltd for suggestions and help in cryo-EM data processing. We also thank the support of Minqi Gao and Xiaoyi Ji from Wuxi Biortus Biosciences Co. Ltd in biochemical assay.

## Author contributions

L.W., L.G. and S.C. conceived the study. J.X., M.W. and X.Y. carried out protein expression and purification, electron microscopy data collection and procession, and structure building and refinement, supervised by S.C. G.Z., D.W. and Z.L. performed functional assays. J.X., L.W., G.Z., D.W., R.F. and S.C. analyzed and interpreted data. J.X., L.W., X.H., R.F. and S.C. wrote the manuscript with input from all authors.

## Extended data figures and tables

**Extended Data Fig. 1.**
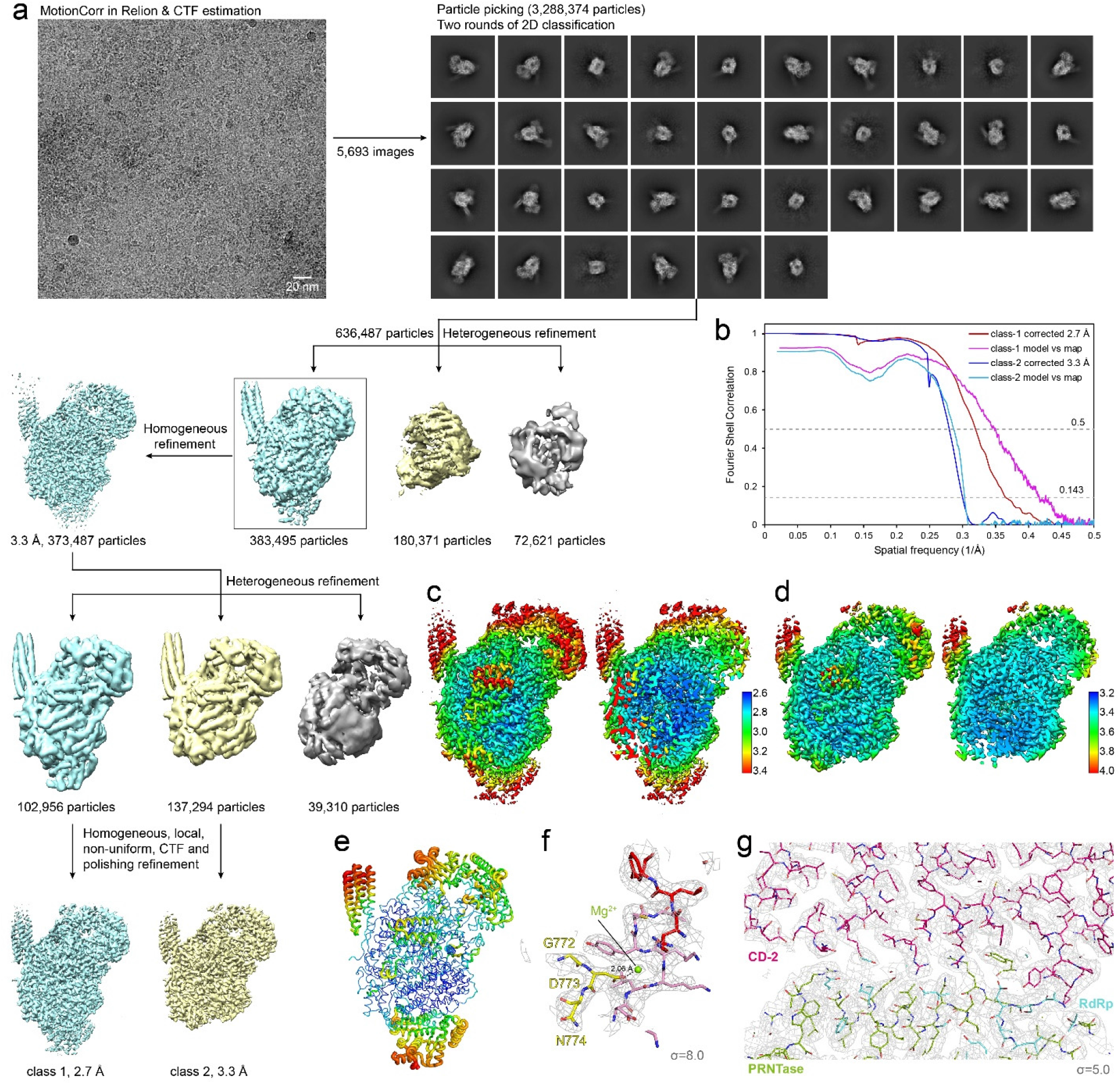
Cryo-EM data processing. **a,** Flow chart of cryo-EM data processing. **b,** The Fourier shell correlation (FSC) curves were used to calculate the resolutions of the final reconstructions with the cutoff of 0.143 and the resolutions are consistent with the model-to-map correlation (0.5 criterion). **c, d,** Local resolution estimation of the cryo-EM density map of class 1 with CD-2 **(c)** and class 2 without CD-2 **(d)**. Overview on the left and cut-out view on the right. **e,** B-factor analysis of the class 1 structure showing the rigidity and flexibility. **f, g,** Representative electron density maps around the RdRp active site including the presumed magnesium ion and catalytic residues 772-GDN-774 **(f),** and at the L-L dimeric interface between CD-2 and RdRp-PRNTase of the intact L **(g)**. The maps are shown as grey mesh, contoured at 8.0 and 5.0 σ, respectively.

**Extended Data Fig. 2.**
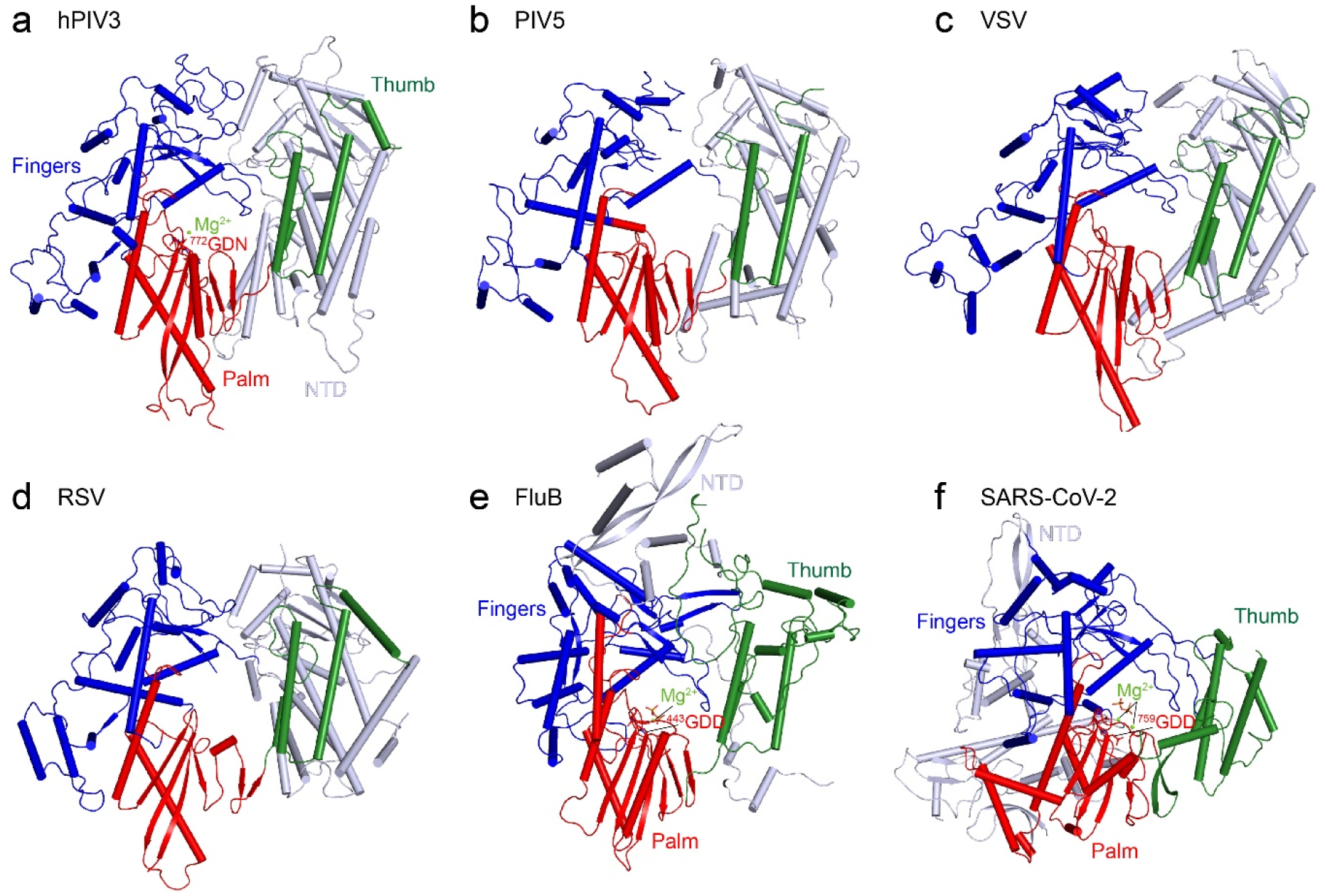
Similarity of the RdRp domains between hPIV3 and other viral polymerases. **a,** hPIV3; **b,** PIV5 (PDB 6V85); **c,** VSV (PDB 6U1X); **d,** RSV (PDB 6PZK); **e,** FluB (PDB 6QCX); **f,** SARS-CoV-2 (PDB 7BV2). The RdRp subdomains are colored as depicted in **Fig. 2a**. The catalytic residues and magnesium ions at the active site in hPIV3, FluB and SARS-CoV-2 RdRps are shown as sticks and sphere, respectively.

**Extended Data Fig. 3.**
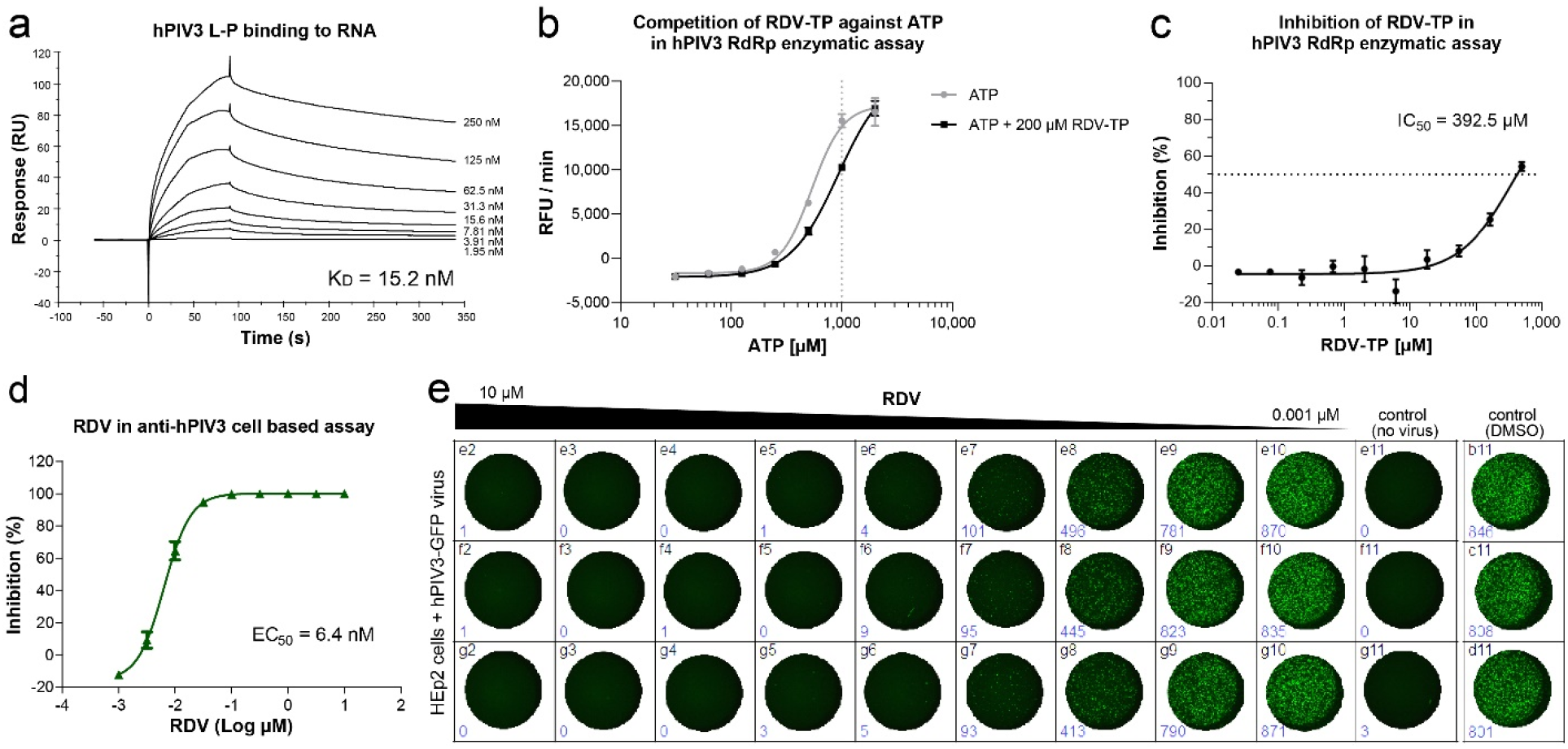
hPIV3 L-P RNA binding activity and the inhibition potency of remdesivir against hPIV3. **a,** The binding affinity of hPIV3 L-P complex to the immobilized RNA duplex was determined by SPR assay. **b,** Enzymatic assay showed that the ATP analog RDV-TP has a competition effect on ATP. **c,** IC_50_ determination of RDV-TP against hPIV3 L-P complex. **d, e,** Fluorescence-based plaque reduction assay was used to determine the EC_50_ of remdesivir (RDV) against hPIV3 in HEp2 cells. All experiments were tested in triplicate.

**Extended Data Fig. 4.**
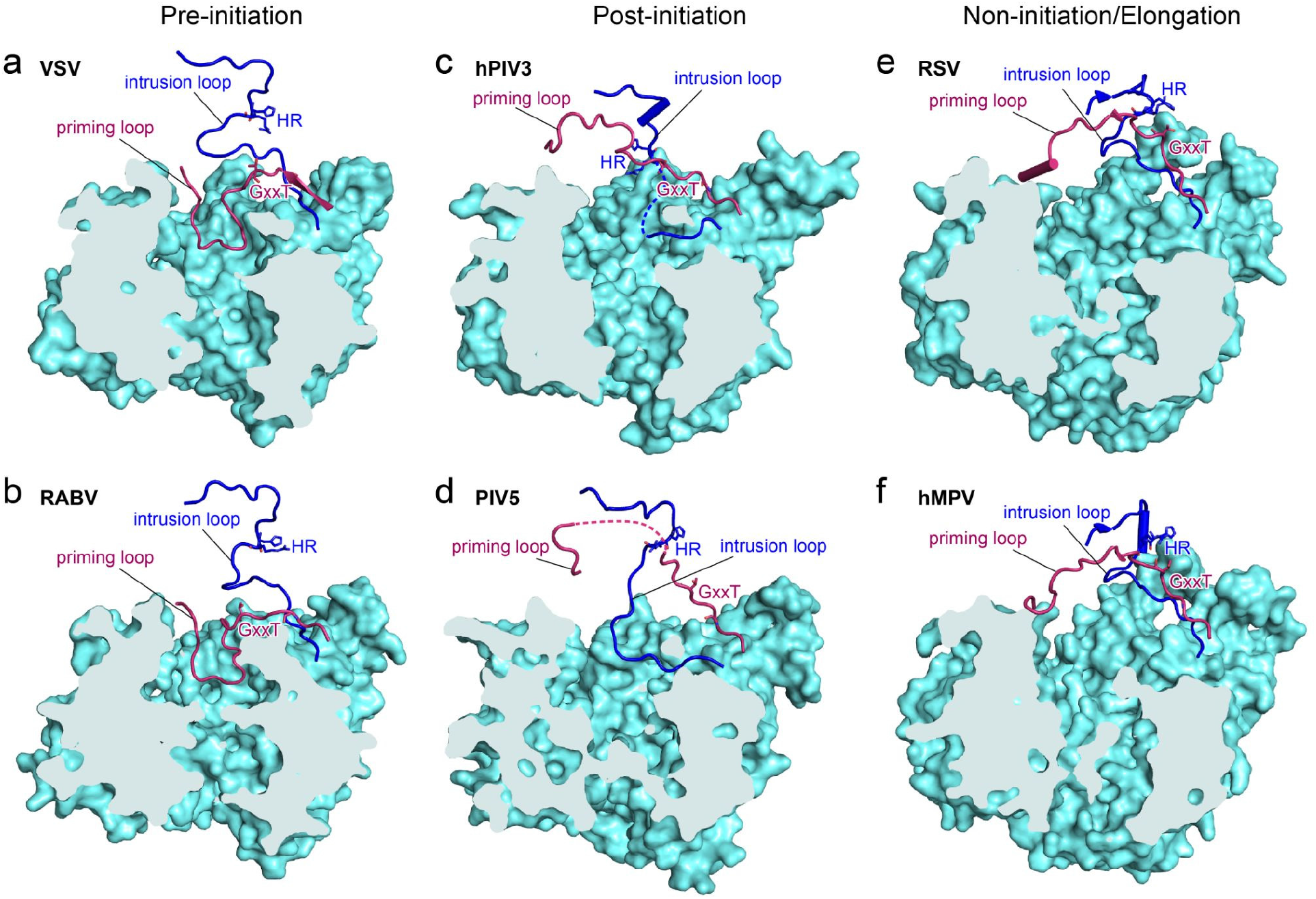
Comparison of the putative priming loops and intrusion loops between hPIV3 and other nsNSV polymerases. **a,** VSV (PDB 6U1X); **b,** RABV (PDB 6UEB); **c,** hPIV3; **d,** PIV5 (PDB 6V85); **e,** RSV (PDB 6PZK); **f,** hMPV (PDB 6U5O). The RdRp domains are shown as aquamarine surface, while the priming and intrusion loops in PRNTase are shown as ribbons in claret and blue, respectively. The conserved GxxT motif and HR motif are shown as sticks. The HR motif is directed towards the RNA cavity in the hPIV3 structure, whereas situated away from the cavity in other structures. The structures adopt different conformations: VSV and RABV, pre-initiation; hPIV3 and PIV5, post-initiation; and RSV and hMPV, non-initiation or elongation.

**Extended Data Fig. 5.**
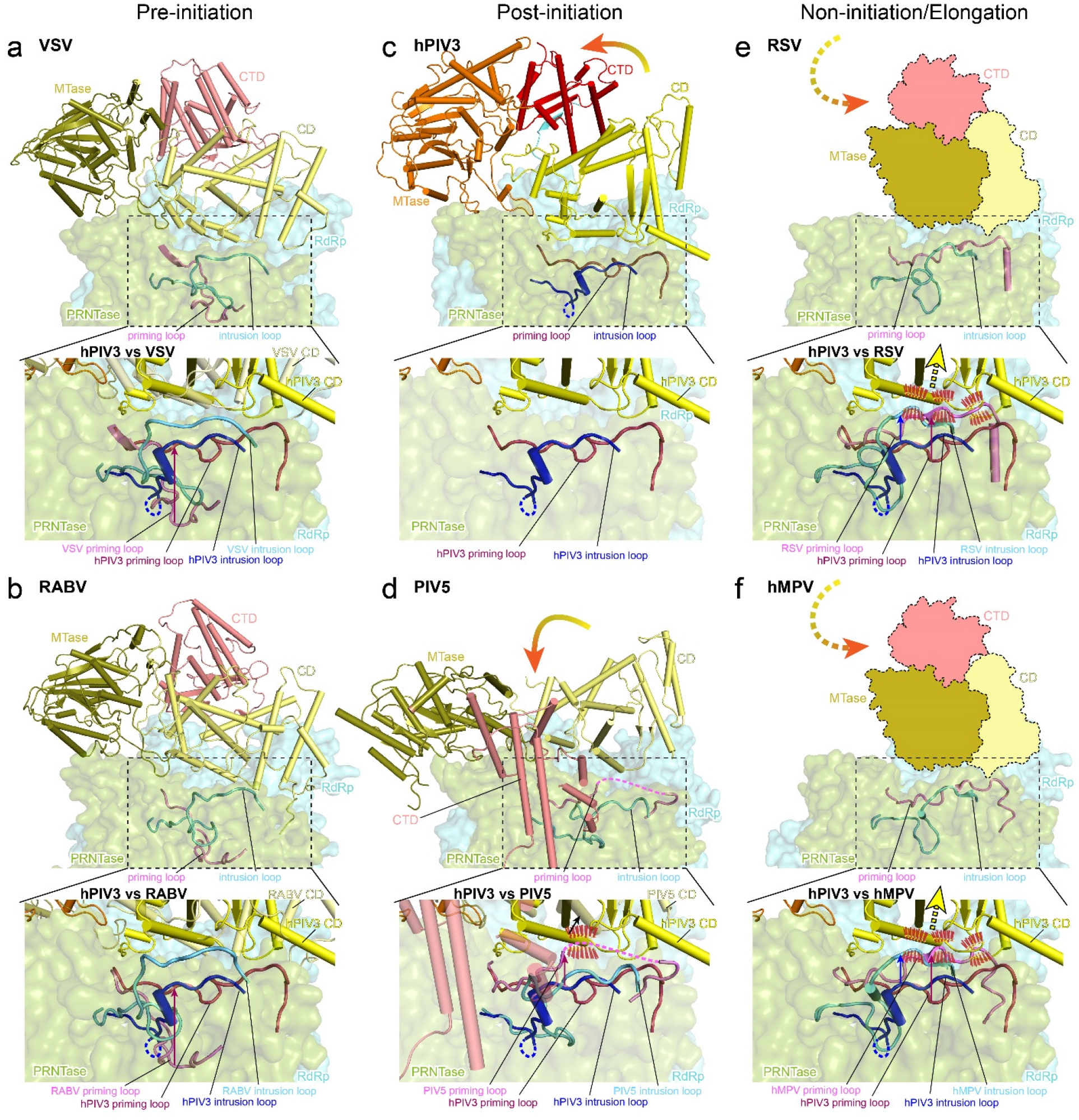
Comparison of L CD-MTase-CTD between VSV. **(a), RABV (b), hPIV3 (c), PIV5 (d), RSV (e) and hMPV (f)** highlights that the interactions between the putative priming/intrusion loops and CD may trigger the movement of CD-MTase-CTD in different states. All the putative priming and intrusion loops and the CD domains are compared to those of hPIV3 and the shifts are indicated by arrows. Potential steric clashes between the priming/intrusion loops of other structures and the CD domain adopting the same conformation as hPIV3 are indicated by red dashes. The movements of CD-MTase-CTD of hPIV3 and PIV5 in post-initiation state compared to that of VSV and RABV in pre-initiation state are indicated by curved arrows. The CD-MTase-CTD that are invisible in VSV and RABV L-P structures are shown as cartoon models with the putative movements shown by dashed arrows. The structures adopt different conformations: VSV and RABV, pre-initiation; hPIV3 and PIV5, post-initiation; and RSV and hMPV, non-initiation or elongation.

**Extended Data Fig. 6.**
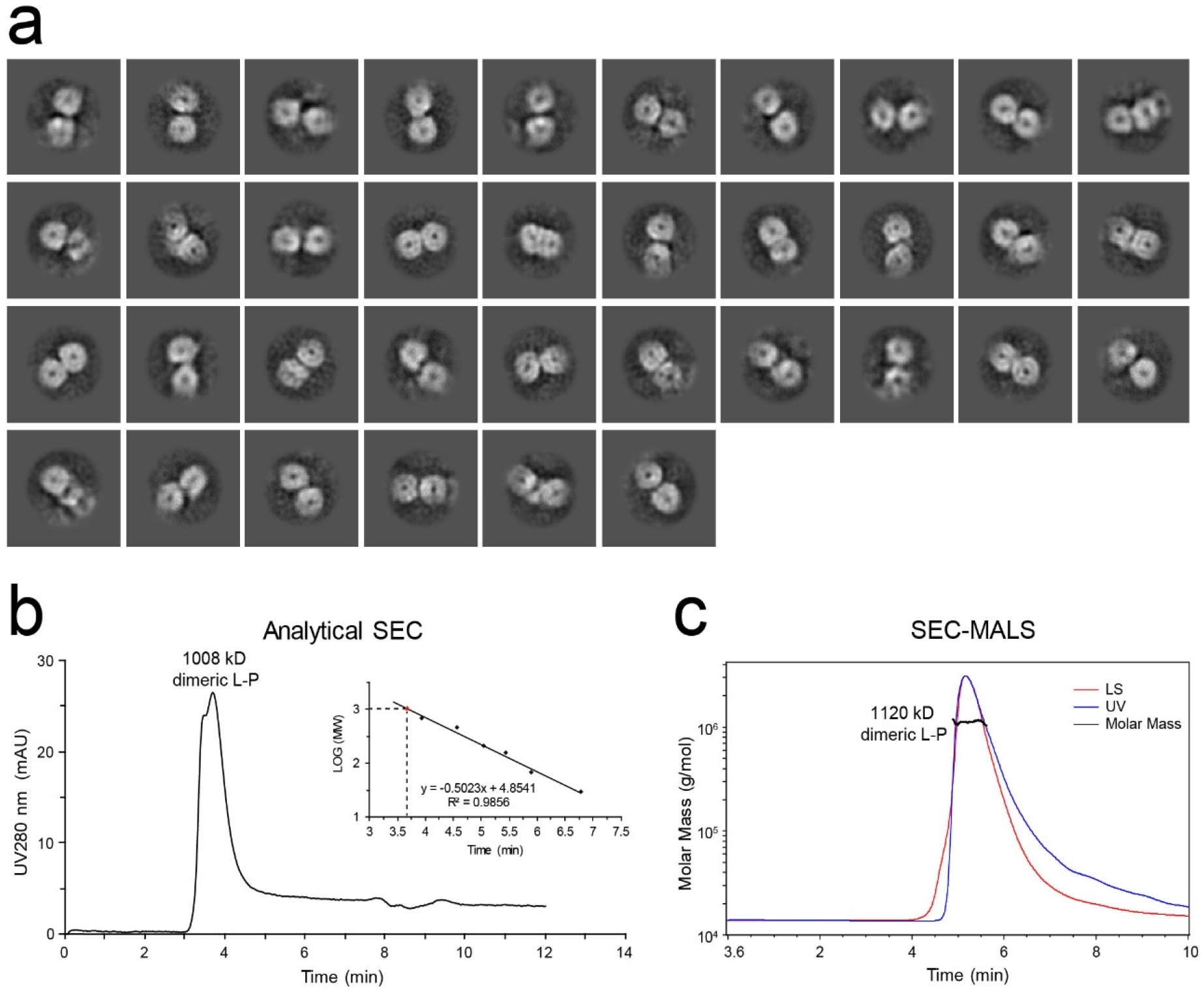
Dimeric L-P polymerase of hPIV3. **a,** 2D classification of the putative hPIV3 L-P dimers. **b,** Molecular weight determination of hPIV3 L-P complex by analytical size-exclusion chromatography. mAU, milliabsorbance units. **c,** Molecular weight determination of hPIV3 L-P complex by size-exclusion chromatography combined with multi-angle light scattering. LS, light scattering; UV, ultraviolet.

**Extended Data Fig. 7.**
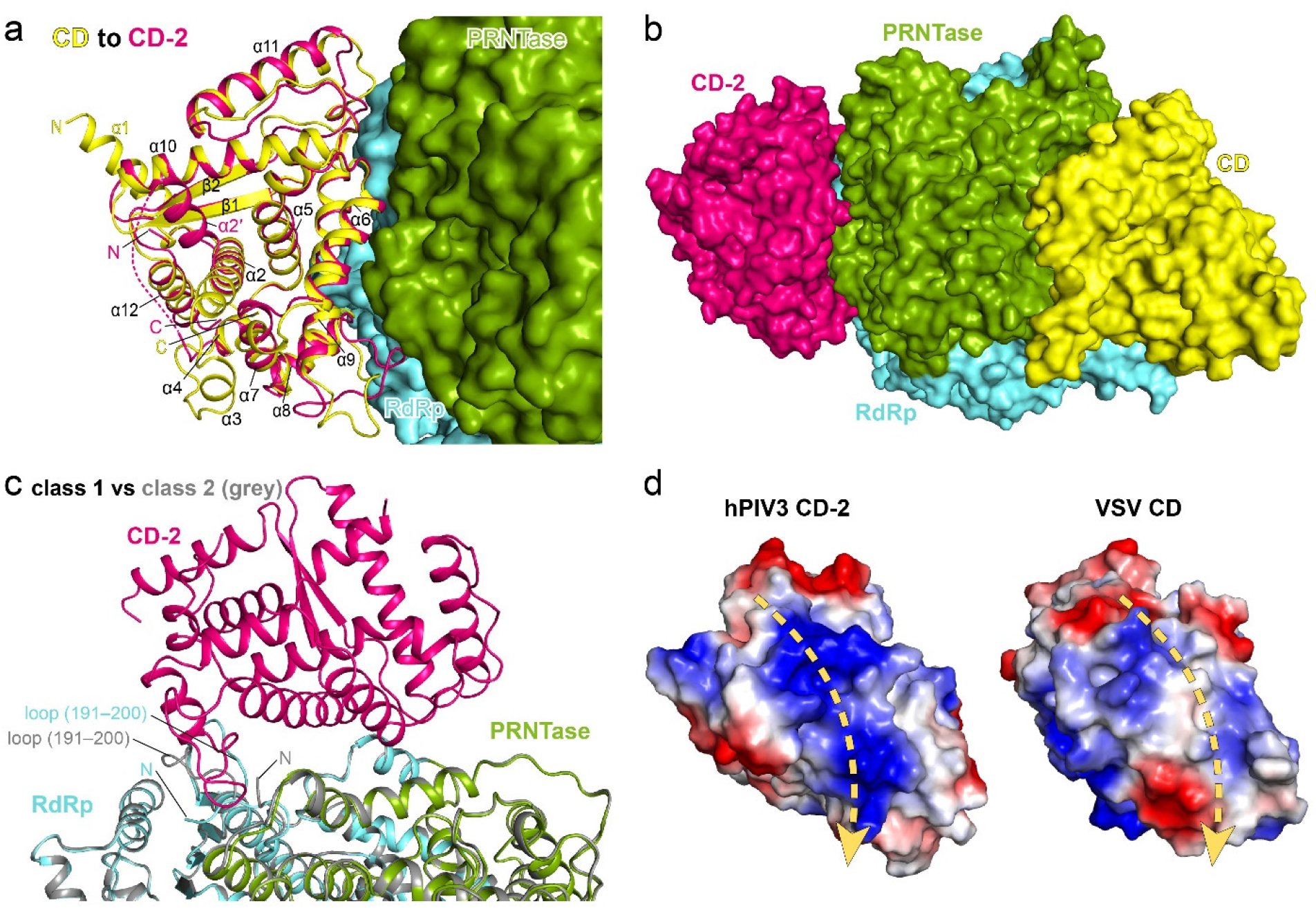
The second CD domain (CD-2) of hPIV3 L. **a,** CD is superimposed to CD-2 with the secondary structure elements and both N and C termini labeled, and RdRp and PRNTase domains are shown as surface. The first α-helix (α1) is invisible in CD-2. The disordered region of CD-2 is shown as a dotted line. The major structural differences are the loops between α8/α9 and α11/β2 at the L-L interface and the bent helix α2 of CD-2. Interestingly, adopting such a bent α2 conformation could cause potential steric clashes for the interface between CD and RdRp-PRNTase within one L. **b,** Two CD domains located at both sides of RdRp-PRNTase of L. All domains are shown as surface. **c,** Comparison of L-P class 1 and class 2 structures. The structural difference of class 2 (the N-terminus and the loop (Asn191–Asp200) of L) is possibly due to the binding of CD-2. **d,** The positively charged surface on CD-2 of hPIV3 L (left) and the comparable surface on the CD domain of the rhabdovirus VSV L (right) at the same top view as in **Fig. 4f**. The putative RNA paths are indicated by dashed arrows.

**Extended Data Fig. 8.**
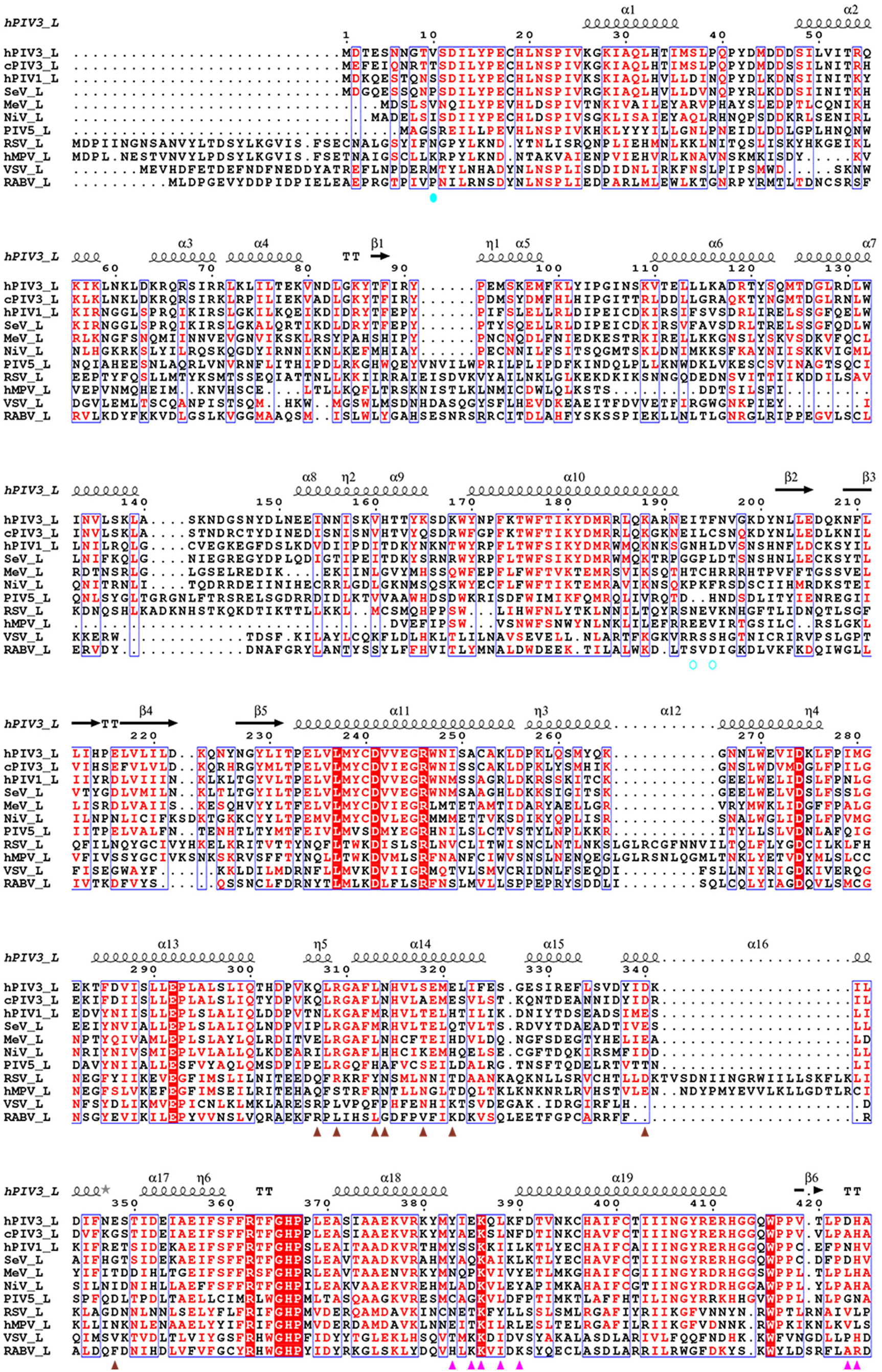
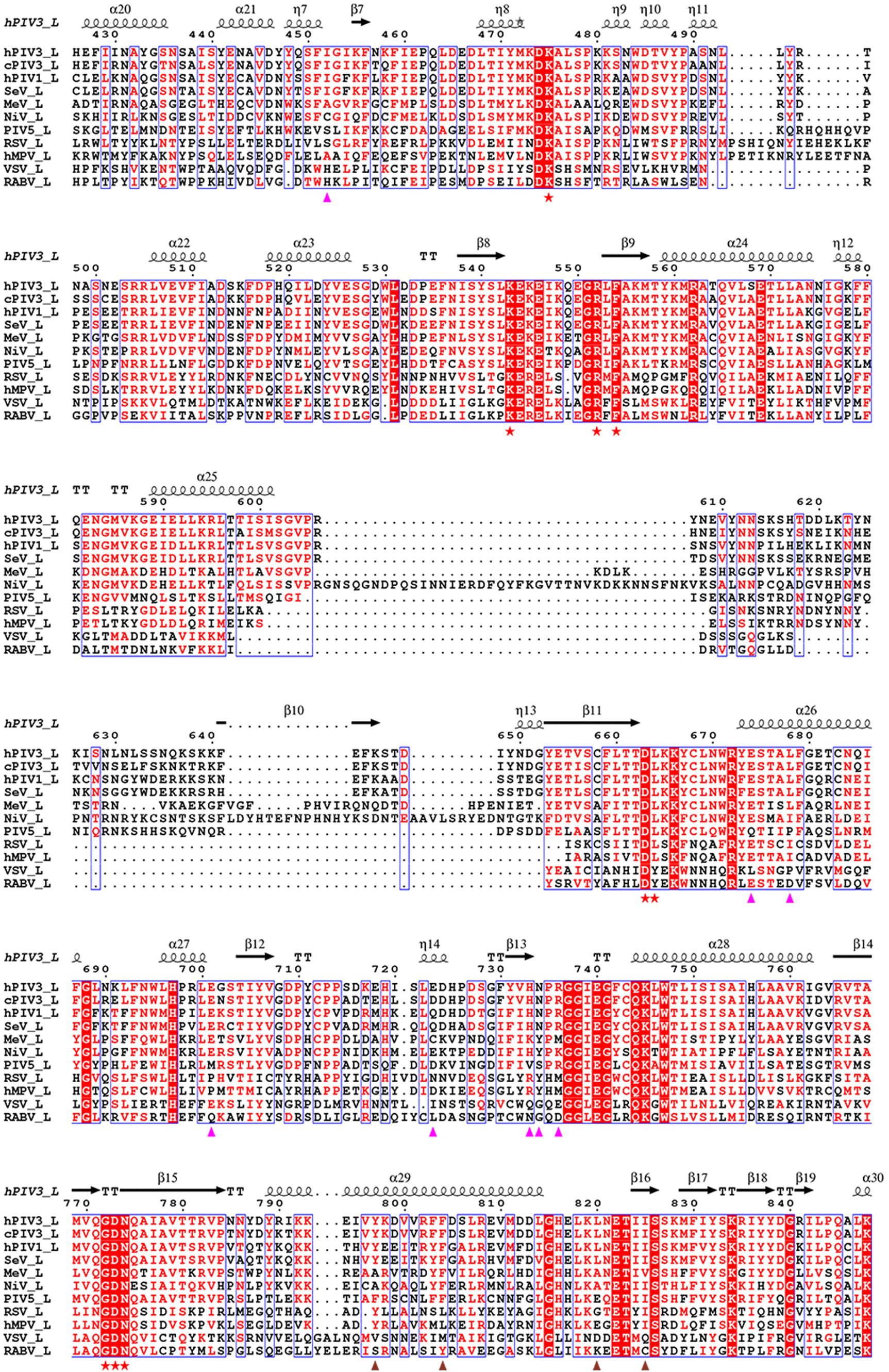
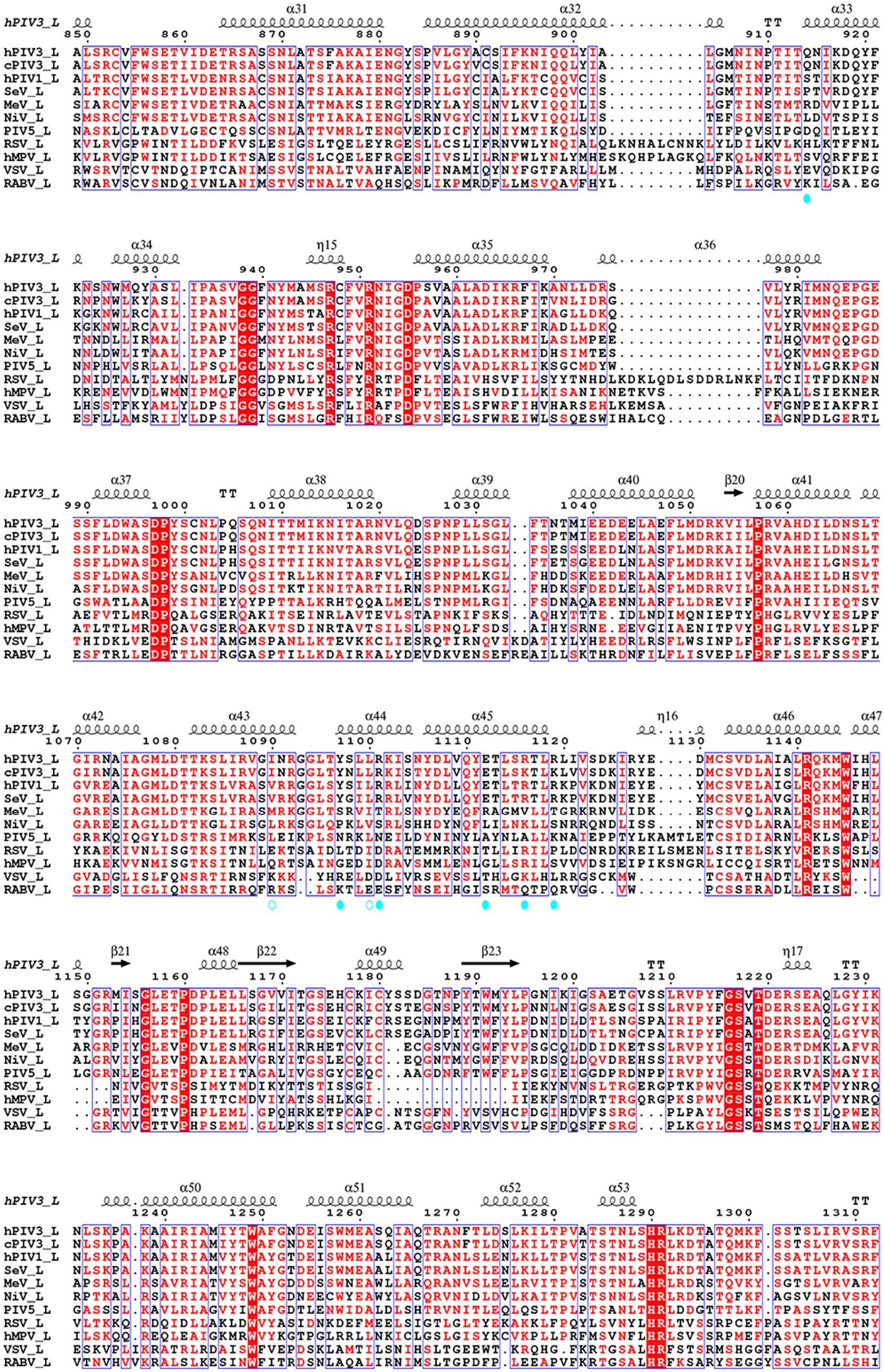
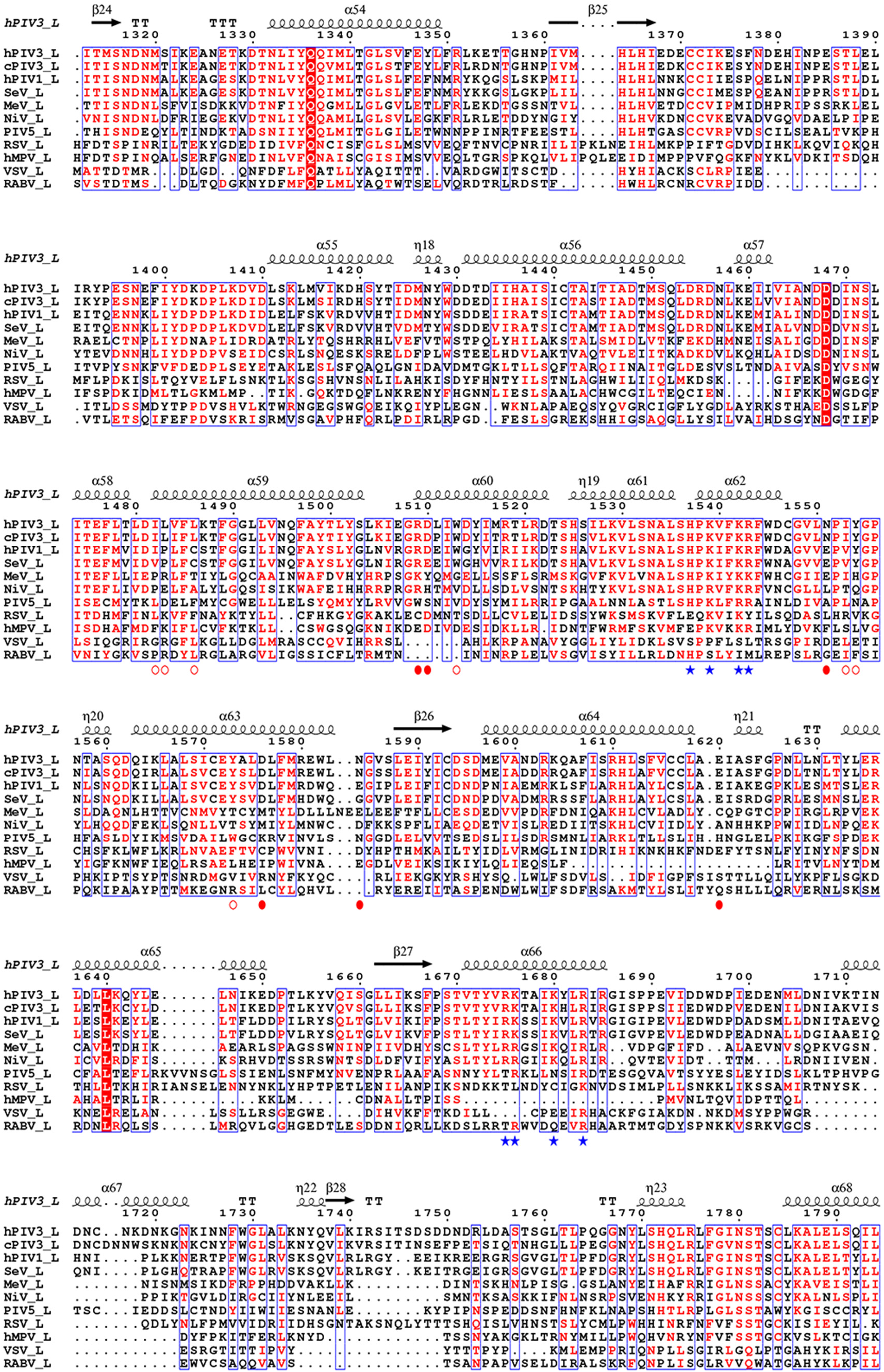
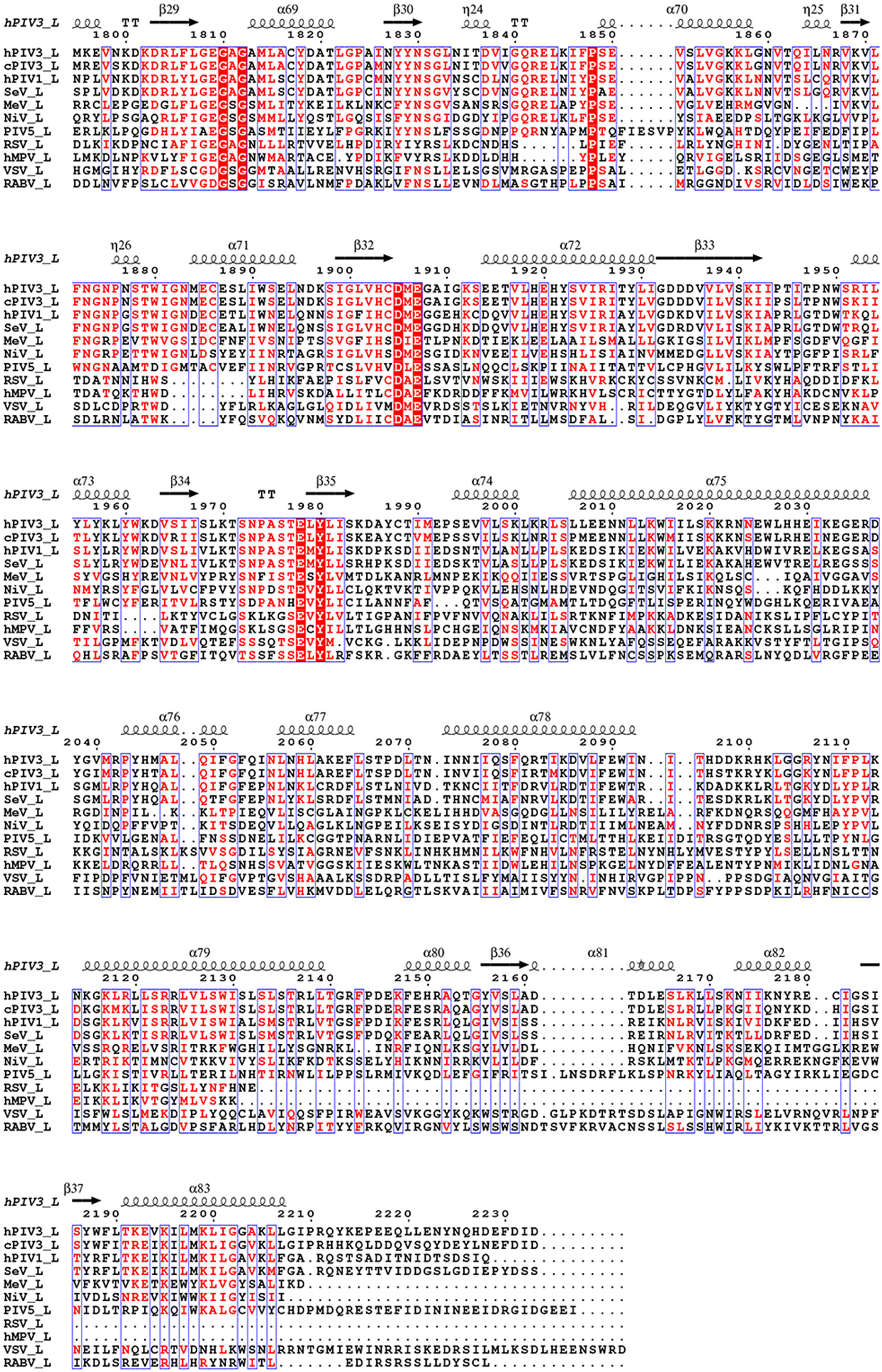
Sequence alignment of nsNSV L proteins. Secondary structure elements of hPIV3 L are shown above the alignment. The presumed catalytic residues and critical residues at the RdRp active site are indicated by red stars below the alignment. The residues involved in L-L interactions of hPIV3 L are indicated by circles: residues on the RdRp and PRNTase domains of the intact L and the CD domain of the second L are colored in cyan and red, respectively; residues participated in polar and hydrophobic interactions are shown as solid and hollow circles, respectively. The basic residues that constitutes the positively charged surface of CD-2 at the putative template RNA entry of the neighboring L are shown as blue stars. The residues of hPIV3 L involved in the interactions with P-OD and P-XD are indicated by pink and maroon triangles, respectively. The sequences of L proteins are from the following viruses: hPIV3, caprine parainfluenza virus 3 (cPIV3, YP_009179212.1), human parainfluenza virus 1 (hPIV1, NP_604442.1), Sendai virus (SeV, BAN84672.1), Measles virus (MeV, NP_056924.1), Nipah virus (NiV, NP_112028.1), parainfluenza virus 5 (PIV5, YP_138518.1), respiratory syncytial virus (RSV, YP_009518860.1), human metapneumovirus (hMPV, Q6WB93), vesicular stomatitis virus (VSV, NP_041716.1) and rabies virus (RABV, ABN11300.1).

**Extended Data Fig. 9.**
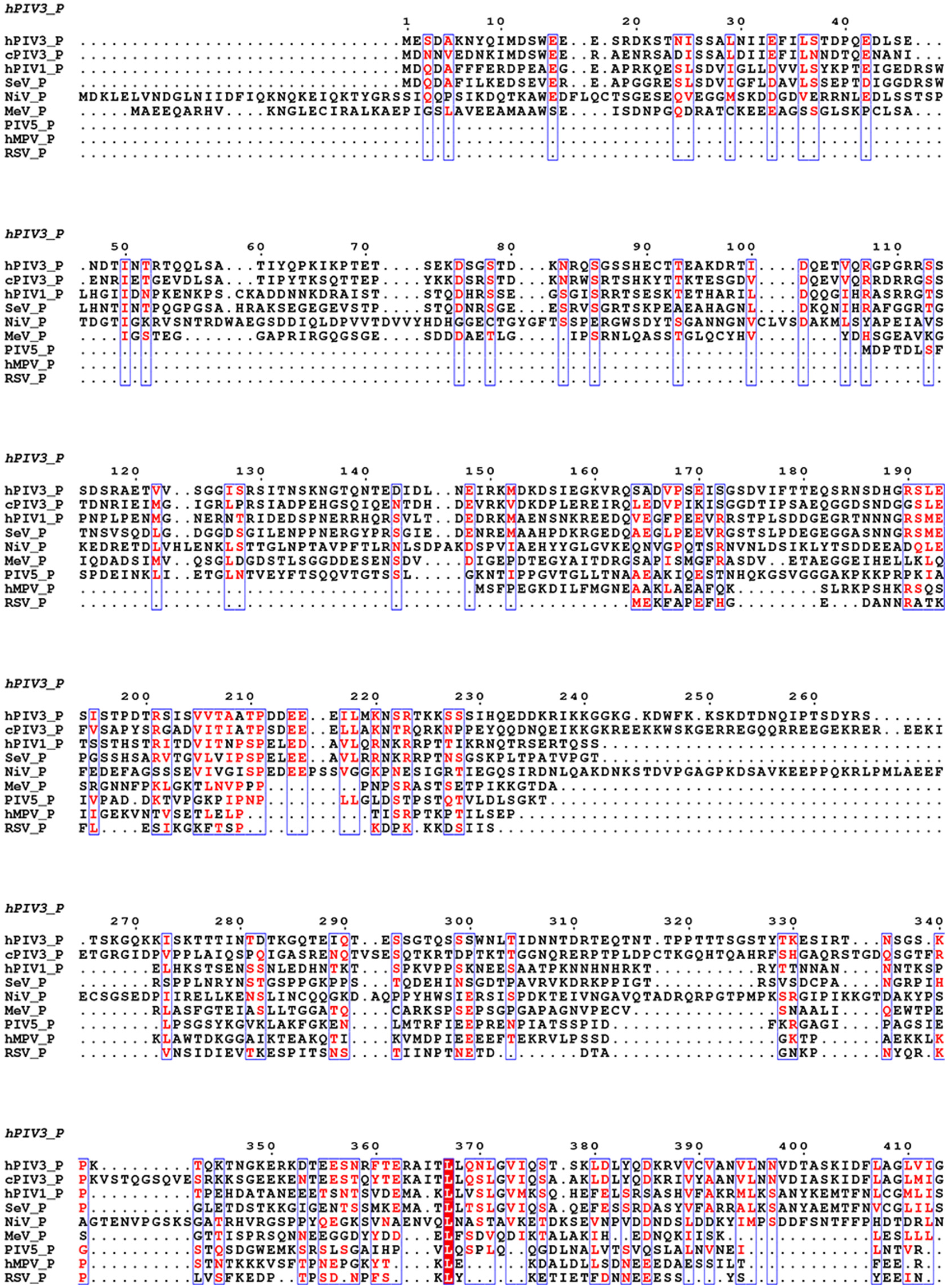
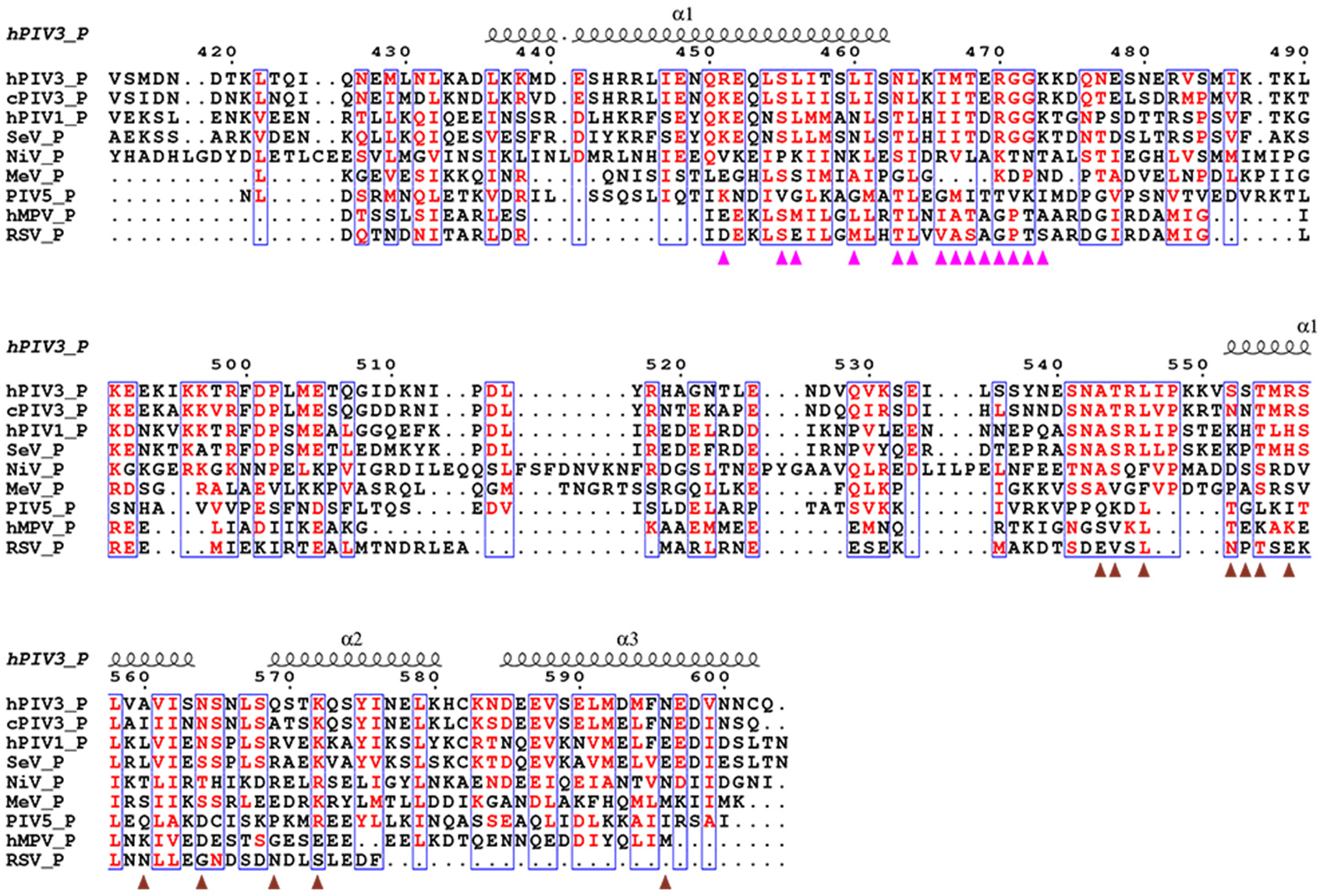
Sequence alignment of nsNSV P proteins. Secondary structure elements of hPIV3 P (P4) are shown above the alignment. The residues of hPIV3 P-OD and P-XD involved in the interactions with L are indicated by pink and maroon triangles, respectively. The sequences of P proteins are from the following viruses: hPIV3, cPIV3 (YP_009179208.1), hPIV1 (NP_604435.1), SeV (BAN84668.1), MeV (NP_056919.1), NiV (NP_112022.1), PIV5 (YP_138512.1), RSV (YP_009518853.1), and hMPV (Q8B9Q8).

**Extended Data Table 1.** Cryo-EM data collection, structure refinement and model statistics.

|  | hPIV3 L:P complex<br>class 1 (dimeric form)<br>(EMD-31755)<br>(PDB 7V70) | hPIV3 L:P complex<br>class 2 (monomeric form)<br>(EMD-31756)<br>(PDB 7V71) |
| --- | --- | --- |
| **Data collection and processing** |  |  |
| Magnification | 81,000 | 81,000 |
| Voltage (kV) | 300 | 300 |
| Electron exposure (e-/Å<sup>2</sup>) | 50 | 50 |
| Defocus range (μm) | -1.5 to -2.0 | -1.5 to -2.0 |
| Pixel size (Å) | 1.087 | 1.087 |
| Symmetry imposed | C1 | C1 |
| Initial particle images (no.) | 3,288,374 | 3,288,374 |
| Final particle images (no.) | 102,956 | 137,294 |
| Map resolution (Å) | 2.7 | 3.3 |
| FSC threshold | 0.143 | 0.143 |
| Map resolution range (Å) | ∞-2.7 | ∞-3.3 |
| **Refinement** |  |  |
| Initial model used (PDB code) | 6V85 | 6V85 |
| Map sharpening *B* factor (Å<sup>2</sup>) | -45 | -109 |
| Model composition |  |  |
| Non-hydrogen atoms | 20,861 | 18,808 |
| Protein residues | 2,584 | 2,329 |
| Ligands | Mg<sup>2+</sup> (1), Zn<sup>2+</sup> (2) | Mg<sup>2+</sup> (1), Zn<sup>2+</sup> (2) |
| *B* factors (Å<sup>2</sup>) |  |  |
| Protein | 70.77 | 92.68 |
| Ligand | 70.00 | 104.78 |
| R.m.s. deviations |  |  |
| Bond lengths (Å) | 0.003 | 0.003 |
| Bond angles (°) | 0.525 | 0.495 |
| Validation |  |  |
| MolProbity score | 1.28 | 1.48 |
| Clashscore | 5.27 | 6.45 |
| Poor rotamers (%) | 0.04 | 0.05 |
| Ramachandran plot |  |  |
| Favored (%) | 98.01 | 97.36 |
| Allowed (%) | 1.99 | 2.64 |
| Disallowed (%) | 0.00 | 0.00 |

